# Combination of NY-ESO-1-TCR-T-cells coengineered to secrete SiRPα decoys with anti-tumor antibodies to augment macrophage phagocytosis

**DOI:** 10.1101/2023.06.27.546523

**Authors:** Evangelos Stefanidis, Aikaterini Semilietof, Julien Pujol, Bili Seijo, Kirsten Scholten, Vincent Zoete, Olivier Michielin, Raphael Sandaltzopoulos, George Coukos, Melita Irving

**Affiliations:** Ludwig Institute for Cancer Research, Department of Oncology, University of Lausanne (UNIL) and University Hospital of Lausanne (CHUV), Lausanne, Switzerland; Department of Molecular Biology and Genetics, Democritus University of Thrace, Alexandroupolis, Greece; Swiss Institute of Bioinformatics, Lausanne, Switzerland; Precision Oncology, University Hospital of Geneva (HUG), Geneva, Switzerland

## Abstract

The adoptive transfer of T cell receptor (TCR)-engineered T cells (ACT) targeting the HLA-A2 restricted cancer-testis epitope NY-ESO-1_157-165_ (A2/NY) has yielded favorable clinical responses against a variety of cancers. Two promising approaches to improve ACT efficacy are TCR affinity-optimization and combinatorial treatment strategies to reprogram the tumor microenvironment (TME). By computational design, we previously developed a panel of affinity-enhanced A2/NY-TCRs. Here, we have demonstrated improved tumor control and engraftment by T cells gene-modified to express one such TCR comprising a single amino acid replacement in CDR3β (A97L). To harness macrophages in the TME, we coengineered TCR-T cells to constitutively or inducibly secrete a high-affinity signal regulatory protein alpha (SiRPα) decoy (CV1) to block the CD47 ‘don’t eat me’ signal. We demonstrated better control of tumor outgrowth by CV1-Fc coengineered TCR-T cells but in subcutaneous xenograft tumor models we observed depletion of both CV1-Fc and CV1 coengineered T cells. Importantly, CV1 coengineered T cells were not depleted by human macrophages in vitro. Moreover, Avelumab and Cetuximab enhanced macrophage-mediated phagocytosis in vitro in the presence of CV1, and augmented tumor control upon ACT. Taken together, our study indicates important clinical promise for harnessing macrophages by combining CV1 coengineered TCR-T cells with tumor-targeting monoclonal antibodies.

## INTRODUCTION

The cancer testis (CT) tumor-associated antigen family member NY-ESO-1 has long been considered a potent candidate for vaccine and immunotherapy development (1). Indeed, NY-ESO-1 is expressed by a broad range of cancers (2, 3) but in healthy adult tissues it is restricted to male germ cells (4). Moreover, NY-ESO-1 has the capacity to elicit spontaneous antibody and T cell responses in a proportion of cancer patients (5, 6). Because NY-ESO-1 is a self-antigen, natural T cell responses tend to comprise TCRs of weaker affinity due to thymic negative selection (7). Notably, TCR-engineered T cells specific for A2/NY, including higher affinity variants (8–10) of the well-characterized TCR 1G4 (11), have demonstrated promising clinical responses against melanoma, synovial cell carcinoma and myeloma (12–16).

Along with TCR affinity-optimization to improve ACT, it is critical to address the TME. On the one hand, the TME can upregulate a range of suppressive mechanisms that must be blocked to unleash the full power of the transferred T cells, and on the other hand there is the potential to harness endogenous immunity against the tumor (17, 18). Indeed, it is now widely held that the eradication of established solid tumors requires the productive interplay of both adaptive and innate immunity (19–21). Reprogramming of the TME can be realized by taking a combinatorial approach, such as the coadministration of radiotherapy (21, 22), chemotherapy, or immune checkpoint blockade (18), for example. Alternatively, the tumor-redirected T cells themselves can be rationally coengineered (23, 24) to constitutively or inducibly express gene-cargo of interest.

CD47, a ‘don’t eat me’ signal recognized by SiRPα on phagocytes, is a cell membrane glycoprotein abundantly expressed by most healthy cells that plays an important role in tissue homeostasis and cell clearance (25). Since the seminal work of Irving Weissman and colleagues (26, 27), numerous studies have reported the upregulation of CD47 as an immune evasion strategy by both liquid and solid cancers (28). In xenograft models (27, 29–32), CD47/SiRPα axis blockade has been associated with enhanced phagocytic activity of macrophages, while in syngeneic models (33–36) dendritic cells (DCs) have emerged as key players, both by augmented phagocytic capacity and their stimulation of adaptive endogenous immunity (36).

However promising, the efficacy of CD47 blockade is hampered by the large CD47 antigen sink in vivo and treatment-related toxicities (37, 38). To address these hurdles, here we sought to take a T-cell coengineering approach to limit blockade of the CD47/SiRPα axis to within the TME. In our study, we evaluated the coengineering of T cells to express a computationally designed A2/NY-TCR (9, 39–41) and to constitutively or inducibly secrete a previously described high-affinity variant of SiRPα, CV1 (38), either as an Fc fusion protein, or as a monomer, and in combination with Cetuximab or/and Avelumab. Taken together, our data indicate important clinical promise for our combinatorial ACT strategy which has the potential to improve tumor control by harnessing both adaptive and innate immunity while lowering the risk of toxicity in patients.

## RESULTS

### Superior tumor control by T cells expressing an affinity-optimized A2/NY TCR

By computational design (39, 40) of an immunodominant A2/NY-TCR isolated from a long-surviving cancer patient (42) we previously developed a panel of increasing-affinity TCRs, one of which has recently shown promise in the clinic (16). The modeling, performed by molecular mechanics generalized Born and surface area solvation (MM-GBSA) free energy calculations (43), was undertaken using the crystal coordinates of the near identical and well-studied TCR 1G4 complexed with A2/NY (11). In previous studies, we demonstrated maximum in vitro effector function (target cell killing, cytokine production, etc.) of T cells gene-engineered to express TCR variants in the upper range of natural affinity (i.e., ∼5-1 μM), beyond which there was an attenuation of activity levels (9, 41), presumably in part due to impaired serial triggering (44). With the aim of improving binding while minimizing the risk of cross-reactivity, one of the TCRs, comprising a single amino acid replacement at position 97 of the CD3β complementary determining loop [A97L; K_D_ = 2.7μM versus 21.4μM for wild-type (WT)], was rationally designed to increase direct interaction with the NY peptide via the CDR3β chain loop (9, 41) (Figure 1A).

**Figure 1.**
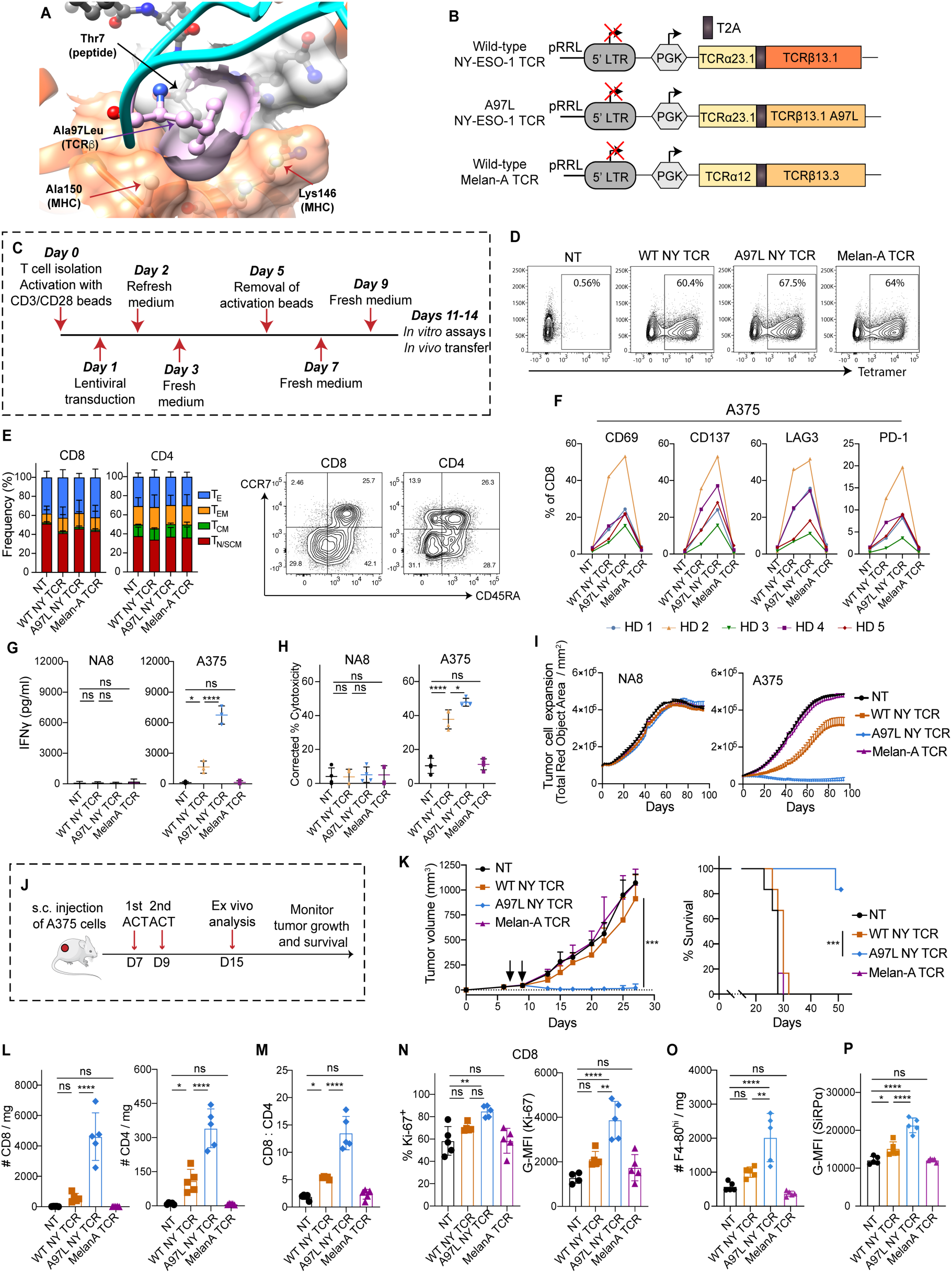
A2/NY-ESO-1 redirected T cells expressing affinity-enhanced TCR A97L exhibit superior effector function and enable significantly improved tumor control and survival as compared to wild type TCR T cells. **A)** TCR variant A97L binding to A2/NY-ESO-1 _157-165_ (PDB ID 2BNR). The amino acid replacement A97L of CDR3β (in ball and stick representation, colored in pink) enhances direct peptide contact via non-polar interactions with Thr7 (in ball and stick representation, colored in grey, below TCRβ A97L) as well as with MHC Ala150 and Lys146 (in ball and stick representation, with surface cored in orange). **(B)** Schematic of lentiviral constructs for expressing TCRs. **(C)** Strategy for T-cell activation, transduction, and expansion. **(D)** Transduction efficiency of CD8^+^ T cells with TCRs evaluated by tetramer staining (data representative of n=5 donors). **(E)** Frequency of effector and memory phenotypes of rested CD8^+^ and CD4^+^ T cells transduced to express the different TCRs (left, n=3). Representative flow cytometric analysis of anti-CCR7 and anti-CD45RA antibody stained T cells (right) (Effector, T_E_; Effector Memory, T_EM_; Central Memory, T_CM_; Naïve/Stem-cell like Memory, T_N/SCM_). **(F)** Expression (frequency) of activation markers and checkpoint receptors on CD8^+^ TCR-engineered T cells 24h post-stimulation with A375 tumor cells (n=5), HD: Healthy Donor. **G)** IFNγ secretion levels by TCR-modified T cells 24h post-stimulation with NA8 and A375 tumor cells at E:T=1:1 (n=3). **H)** Frequency of AnnexinV^+^ DAPI^+^ cells, corrected to tumor alone, in 24h co-cultures of NA8 and A375 tumor cells with TCR-modified T cells at E:T = 1:1 (n=4). **I)** Evaluation of mKate2^+^ NA8 and A375 tumor cell growth control over time by TCR-modified T cells at E:T = 1:1 using live-cell IncuCyte imaging (data representative of n=4 donors). **J)** Schematic of ACT study. **K) C**ontrol of A375 tumors (left) in NSG mice and survival curves (right) following adoptive transfer of TCR-modified T cells (n=6 mice per group, data representative of 2 independent studies). **L)** Number of intratumoral human CD8^+^ (left) and CD4^+^ (right) T cells per mg of tumor 7 days post-ACT (n=5, data representative of 2 independent studies). **M)** Ratio of intratumoral CD8^+^:CD4^+^ human T cell frequency 7 days post-ACT (n=5, data representative of 2 independent studies). **N)** Frequency (left) and G-MFI (right) of Ki-67 expression within intratumoral human CD8^+^ T cells 7 days post-ACT (n≥4, data representative of 2 independent studies). **Ο**) Number of intratumoral mouse F4-80^hi^ macrophages per mg of tumor 7 days post-ACT (n=5, data representative of 2 independent studies). P) G-MFI of mouse SiRPα expression within intratumoral mouse F4-80^hi^ macrophages 7 days post-ACT (n=5, data representative of 2 independent studies). Statistical analysis by one-way analysis of variance (ANOVA) (E, G-H, L-P), two-way ANOVA (K, left) or Mantel-Cox (K, right) with correction for multiple comparisons by post hoc Tukey’s test (E, G-H, K-P). ****P< 0.0001; ***P < 0.001; **P < 0.01; *P < 0.05.

We began by subcloning the WT- and A97L-TCR into a lentiviral transfer vector (Figure 1B). As a negative control, we similarly subcloned an HLA-A2 /Melan-A_57-65_-directed TCR (Lau444; Melan-A TCR) derived from a long-surviving cancer patient (45). Primary human CD4^+^ and CD8^+^ T cells were subsequently transduced with TCR-encoding lentiviruses and expanded (Figure 1C). Tetramer staining of transduced CD8^+^ T cells revealed similar high cell-surface expression levels of the TCRs (Figure 1D). Because the WT A2/NY TCR is CD8-dependent (9), we used an anti-Vβ13.1 mAb to detect its expression by gene-modified CD4^+^ T cells (Supplemental Figure 1A). We observed similar expansion (Supplemental Figure 1B), as well a proportion of effector (T_E_), effector memory (T_EM_), central memory (T_CM_) and naïve/stem-cell like (T_N/SCM_) phenotypes (Figure 1E left) as assessed by cell-surface expression of CCR7 and CD45RA for the different TCR-engineered CD8^+^ and CD4^+^ T cells by flow cytometry (Figure 1E right, Supplemental Figure 1C).

We subsequently evaluated the TCR engineered T cells both in vitro and in vivo. Co-culture with the A2^+^/NY^+^ melanoma cell line A375 revealed consistently higher upregulation of the activation markers CD69 and CD137 as well as of the inhibitory markers LAG3 and PD-1 amongst donors by CD8^+^ A97L-TCR T cells as compared to WT-TCR ones (Figure 1F, Figure S1D). Moreover, upon co-culture with A375, we observed significantly higher IFNγ production and cytotoxicity by A97L-TCR T cells than WT-TCR ones (Figure 1, G and H). Similar results were acquired using the A2^+^/NY^+^ osteosarcoma cell line Saos-2 as T-cell target (Supplemental Figure 1, E and 1F). Likewise, in an IncuCyte assay, in which tumor cell control by T cells is tracked by live cell imaging over days, A97L-TCR T cells were far more potent than WT ones against A375 (Figure 1I). Importantly, there was no reactivity of the WT-nor A97L-TCR T cells against the A2^+^/NY^-^ melanoma tumor cell line NA8, nor did MelanA-TCR T cells show any reactivity above background levels (i.e., of non-transduced (NT) T cells) in any of the assays against NA8, A375, and Saos-2 (Figure 1, F-I, Supplemental Figure 1, D-G).

Next, we performed ACT studies (using CD8^+^:CD4^+^ T cells at a 4:1 ratio) in NSG mice bearing subcutaneous A375 tumors (Figure 1J) and observed significant tumor control and improved survival (Figure 1K) upon treatment with A97L-TCR T cells but not for WT-TCR-T cells nor the control T cells. Ex vivo characterization at day 7 post-ACT revealed significantly higher intratumoral presence of both CD8^+^ and CD4^+^ A97L-TCR-T cells than WT-TCR ones (Figure 1L), as well as a significantly higher CD8:CD4 ratio for A97L-TCR-T cells (Figure 1M). In addition, the CD8^+^ A97L-TCR TILs were characterized by higher Ki-67 levels (both % and MFI, Figure 1N), indicative of superior proliferative capacity of the affinity-optimized TCR-T cells. Despite the elevated numbers of CD4^+^ A97L-TCR-T cells in the TME (Figure 1L), there were no differences in Ki-67 expression (Supplemental Figure 1H) suggesting either superior homing or/and persistence (survival and retention) in the TME. Notably, ACT with A97L-TCR-T cells was accompanied by a significantly increased presence of intratumoral F4-80^hi^ mouse macrophages (Figure 1O) that expressed higher levels of the inhibitory receptor SiRPα (Figure 1P).

Thus, taken together we demonstrated significantly higher in vitro effector function and tumor control by T cells bearing the affinity optimized A97L-TCR as compared to the WT-TCR, the latter of which appears to be supported by superior proliferative capacity of the A97L-TCR-T cells. However, the elevated levels of SiRPα^hi^ F4-80^hi^ mouse macrophages in the tumors of A97L-TCR-T cell treated mice may counteract tumor control. Therefore, we next sought an approach of safely targeting the SiRPα/CD47 axis directly within the TME by means of coengineered TCR-T-cells.

### CV1 SiRPα-Fc promotes macrophage-mediated phagocytosis of tumor cells

We considered two T-cell coengineering approaches to target the SiRPα/CD47 axis; first, the local secretion of a single chain variable fragment (scFv; specific for CD47) as has previously been demonstrated effective in targeting PD-1, for example (46), and the second, coengineering with a previously developed variant of the SiRPα ectodomain, CV1 (38). Notably, CV1 is of high affinity for CD47, small in size, and has demonstrated low toxicity and short half-life upon systemic administration as compared to antibody-based inhibitors of CD47 (38). These are all favorable properties for gene-cargo that will be delivered and replenished within the TME by tumor-specific T cells. We thus set out to coengineer A97L-TCR T cells with CV1.

We began by validating the binding and functionality of soluble recombinant proteins comprising the CD47-binding ectodomain of human SiRPα (amino acids 27-371) fused to human IgG1-Fc which has been shown to enhance tumor cell phagocytosis by macrophages (47). Throughout our study, we compare the WT sequence with the previously described high-affinity variant CV1 (38) as well as an inactive (in) (48) variant of the SiRPα ectodomain (Figure 2A). The A2^+^/NY^+^ tumor cell lines Me275, A375 and Saos-2 express CD47 at their surface (Figure 2B top) and as expected we observed higher binding (Figure 2B bottom), as well as saturation of binding at lower concentrations (Figure 2C), for CV1 SiRPα-Fc than the wtSiRPα-Fc recombinant fusion protein. As expected, there was no binding to CD47 by inSiRPα-Fc (Figure 2B bottom, Figure 2C), nor was there binding by any of the fusion proteins to CD47-deficient JinB8 Jurkat cells (Figure 2D), hence validating specificity of CV1 for CD47.

**Figure 2.**
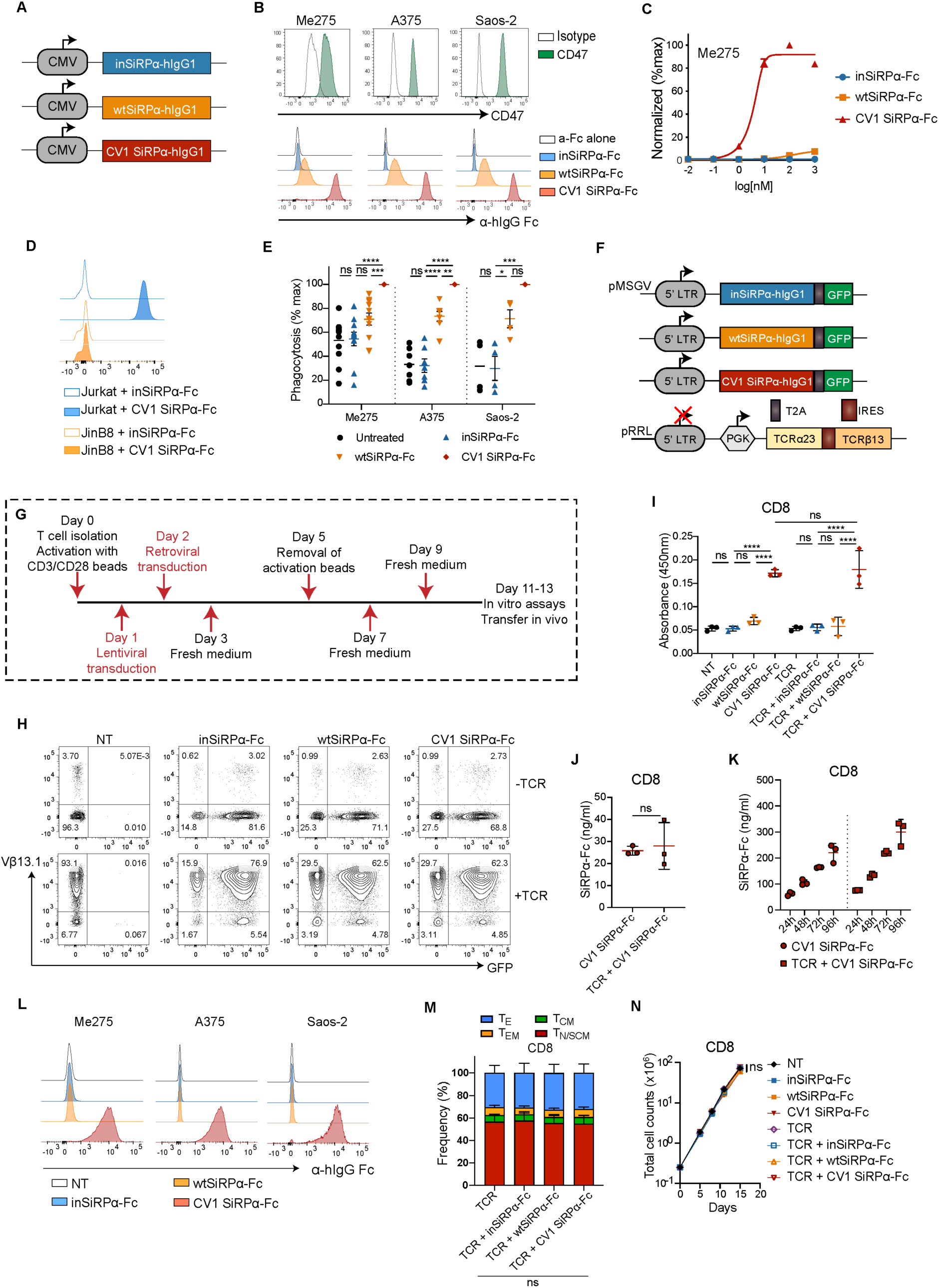
Soluble SiRPα-Fc decoys bind tumor-expressed CD47 enabling enhanced phagocytosis by human MDMs, and they can be efficiently secreted by gene-modified T cells. **A)** Schematic of expression vectors encoding human inSiRPα-Fc, wtSiRPα-Fc and CV1 SiRPα-Fc proteins. **B)** Panel of HLA-A2^+^ NY-ESO-1^+^ tumor cell lines expressing CD47 on their cell surface (top) and detection of wtSiRPα-Fc and CV1 SiRPα-Fc soluble protein binding on tumor CD47 by anti-human IgG Fc mAb staining (bottom). **C)** Binding of increasing concentration of SiRPα-Fc to CD47 on Me275 tumor cells (data representative of n=2 independent studies). **D)** SiRPα-Fc binding profile on CD47^+^ Jurkat and CD47-deficient JinB8 cells (data representative of n=3 independent studies). **E)** Phagocytosis of PKH26-labeled tumor cells by human MDMs in the presence of 10ug/ml soluble SiRPα-Fc proteins (inactive versus CV1). Phagocytosis is normalized to the maximal response of each individual donor (n ≥ 4). Data represent mean ± SEM. **F)** Schematic of retroviral constructs encoding inSiRPα-Fc, wtSiRPα-Fc and CV1 SiRPα-Fc proteins followed by an eGFP reporter gene, and of a lentiviral construct encoding the A97L NY-ESO-1 TCR. **G)** Strategy for T-cell activation, dual virus transduction and expansion. **H)** Expression of SiRPα-Fc molecules and A97L TCR in co-transduced rested human CD8^+^ T cells as detected by eGFP and anti-human Vβ13.1 mAb, respectively (data representative of n=12 independent donors). **I)** CD47-based ELISA detection of SiRPα-Fc secreted by engineered CD8^+^ T cells (n=3). **J)** Quantification of CD8^+^ T cell-secreted CV1 SiRPα-Fc by CD47-based ELISA (n=3). **K)** Quantification of CV1 SiRPα-Fc accumulated in culture supernatants of engineered CD8^+^ T cells over time (representative results for n=3 donors). **(L)** Binding of CD8^+^ T cell-secreted high-affinity CV1-SiRPα-Fc on different CD47^+^ tumor cell lines (data representative of n=3 donors). **M)** Frequency of effector and memory phenotypes of transduced and rested CD8^+^ T cells (n=3) (Effector, T_E_; Effector Memory, T_EM_; Central Memory, T_CM_; Naïve/Stem-cell like Memory, T_N/SCM_). **N)** Expansion of engineered CD8^+^ T cells (n=3). Statistical analysis by one-way analysis of variance (ANOVA) (E, I and M), unpaired two-tailed t test (J), or two-way ANOVA (N) with correction for multiple comparisons by post hoc Tukey’s test on pooled donors (E, I-J and M-N). ****P< 0.0001;***P < 0.001; **P < 0.01; *P < 0.05.

To compare the ability of the SiRPα-Fc molecules to promote phagocytosis, we set up co-cultures of human CD14^+^ monocyte-derived macrophages (MDMs) with PKH26 fluorescently pre-labeled tumor targets and the cells were analyzed 4 hours later by flow cytometry (Figure 2E; gating demonstrated in Supplemental Figure 2A) and Amnis imaging (Supplemental Figure 2B). Consistent with binding properties, CV1 SiRPα-Fc promoted a significantly higher level of tumor cell phagocytosis by macrophages than WT, and the inactive variant had no biological impact (Figure 2E).

Encouraged by our results, we moved forward with building retroviral cassettes encoding CV1-, WT- and inSiRPα-Fc molecules along with GFP to facilitate detection of transduced T cells (Figure 2F). Subsequently, we co-transduced activated primary human CD8^+^ and CD4^+^ T cells with lentivirus for the A2/NY TCR and 24 hours later with retrovirus encoding the SiRPα-Fc molecules (Figure 2G). The use of lenti- and retroviruses purposefully packaged with VSV-G and RD114 (49), respectively, was undertaken to abrogate virus envelope glycoprotein competition for target receptors enabling T cell infection. Our mixed virus strategy led to high dual transduction efficiency in both CD8^+^ (>60%) and CD4^+^ (>50%) T cells, as evaluated by simultaneous detection of GFP, as a surrogate for the decoys, and the A2/NY-TCR by Vβ13.1 staining (Figure 2H, Supplemental Figure 2C).

A CD47-based ELISA confirmed the secretion of the different SiRPα-Fc molecules by gene-modified CD8^+^ and CD4^+^ T cells (Figure 2I, Supplemental Figure 2D), and that similar quantities of CV1 SiRPα-Fc were produced upon co-transduction of the T cells to express the A97L-TCR (Figure 2J, Supplemental Figure 2E). Furthermore, we observed an accumulation of CV1 SiRPα-Fc in the culture supernatant of the gene-modified T cells over time, reaching approximately 300ng/mL for CD8^+^ A97L-TCR T cells (Figure 2K) and 450ng/mL for CD4^+^ A97L TCR-T cells (Supplemental Figure 2F) at 96 hours. Notably, this is almost an order of magnitude lower than the working concentration typically used for CD47-blocking monoclonal antibodies (mAbs) and recombinant SiRPα-Fc proteins (38) and likely explains the detection of T-cell secreted CV1-but not WT SiRPα-Fc binding to CD47-expressing tumor cells (Figure 2L, Supplemental Figure 2G) from culture supernatants.

Finally, we tested if the dual viral transduction had any impact on the biological properties of engineered T cells. We observed no differences in memory phenotype (Figure 2Μ, Supplemental Figure 2H) nor in expansion (Figure 2N, Supplemental Figure 2I) between the different single (decoy or TCR alone) or dual (TCR plus decoy) engineered T cells versus NT T cells.

Hence, in summary, and consistent with previous reports (38), we demonstrated higher binding of CV1 SiRPα-Fc than wtSiRPα-Fc to tumor-cell surface expressed CD47, as well as superior phagocytosis of tumor cells by human MDMs in the presence of CV1 SiRPα-Fc. Furthermore, dual transduction with lenti- and retrovirus enabled high coexpression in T cells of A97L-TCR and CV1 SiRPα-Fc decoys, the latter of which were secreted into the supernatant and shown to bind tumor cell surface expressed CD47.

### CV1 SiRPα-Fc secreted by TCR-T cells augments macrophage-mediated phagocytosis

We next sought to evaluate the ability of SiRPα-Fc decoy secreted by gene-modified T cells to enhance macrophage-mediated phagocytosis of tumor cells. First, however, we tested if there was any direct impact of the decoys on T-cell function in the presence of target tumor cells (schematic in Supplemental Figure 3A). While there were no differences in specific target-cell killing by TCR-T cells coengineered or not with the decoy (Supplemental Figure 3, B-D), we consistently observed higher IFNγ production amongst donors by CV1 SiRPα-Fc coengineered TCR-T cells (Supplemental Figure 3E), in line with previous work demonstrating a co-stimulatory role for CD47 on T cells (50).

We generated human CD14^+^ MDMs in order to evaluate the ability of T-cell secreted SiRPα-Fc decoys to promote phagocytic activity against tumor cells. To this end, co-cultures of tumor cells and MDMs were performed in the presence of supernatants collected from decoy-engineered T-cell cultures (Figure 3A). We noted significantly higher phagocytic activity of MDMs against tumor cells in the presence of CV1-than wtSiRPα-Fc which, like inSiRPα-Fc, and consistent with its lack of binding at secreted concentrations (Figure 2L, Supplemental Figure 2G), had no effect on phagocytosis (Figure 3B). Supernatants from the culture of CV1 SiRPα-Fc T cells had a similar impact on phagocytosis regardless of whether or not the T cells were coengineered to express the A97L-TCR (Supplemental Figure 3F), in line with the fact that similar amounts of the high-affinity decoy are produced with or without the TCR. Finally, we set up triple co-culture assays comprising target tumor cells, MDMs and the differently engineered T cells (schematic in Figure 3C) and consistently observed, even in a just a 4-hour period (i.e., limited time for decoy secretion into the culture supernatant), higher phagocytic activity in the presence of CV1 SiRPα-Fc-engineered T cells than ones expressing inSiRPα-Fc (Figure 3D).

**Figure 3.**
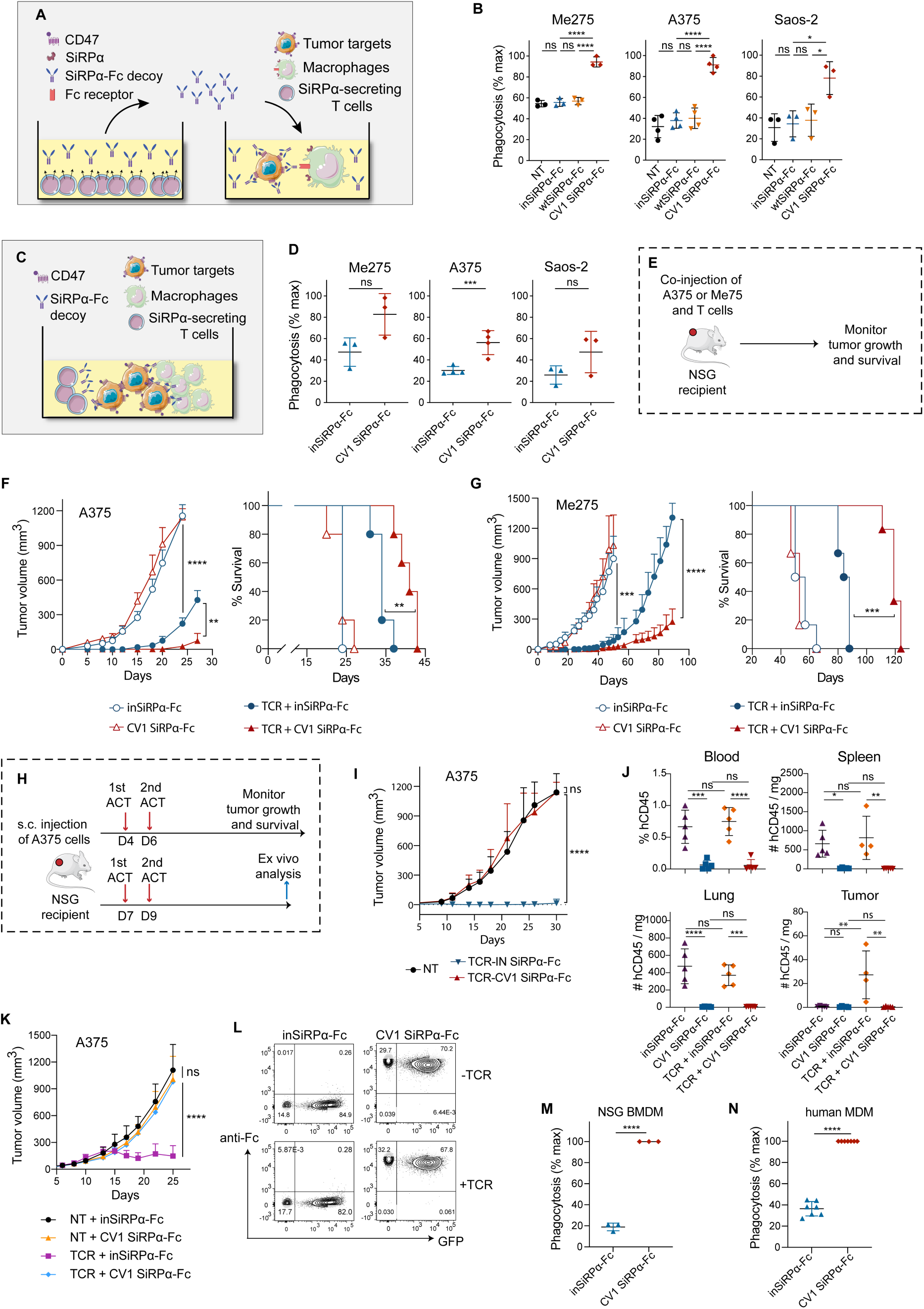
T-cell secreted CV1 SiRPα-Fc decoys augment tumor cell phagocytosis by macrophages in vitro. **(A)** Schematic of tumor cell phagocytosis by macrophages (human MDMs) in the presence of supernatant from SiRPα-Fc-engineered T cells. **(B)** Tumor cell phagocytosis by MDMs in the presence of supernatant from SiRPα-Fc-engineered T cells (n≥3). **(C)** Schematic of triple co-culture of tumor cells, macrophages (MDMs) and engineered T cells. **(D)** Tumor cell phagocytosis by MDMs in triple co-cultures with SiRPα-Fc-secreting T cells (n≥3). **(E)** Schematic of Winn assay. **(F)** Control of A375 outgrowth (left) and survival (right) in a Winn assay with SiRPα decoy co-engineered A97L-TCR T cells (n=5, data representative of n=2 independent studies). **(G)** Control of Me275 outgrowth (left) and survival (right) in a Winn assay with SiRPα decoy co-engineered A97L-TCR T cells (n=6, data representative of n=2 independent studies). **(H)** Schematic of ACT against subcutaneous A375 tumors and ex vivo analysis. (**I**) A375 tumor growth and control curves following ACT (n=7). (**J**) Frequency and number of human CD45^+^ cells in harvested tissues 5 days post-ACT (n≥4, data representative of n=2 independent studies). (**K**) A375 tumor growth and control curves following ACT supplemented with co-administration of soluble inSiRPα-Fc and CV1 SiRPα-Fc proteins (n≥5). (**L**) Flow cytometry detection of T cell-secreted CV1 SiRPα-Fc binding on T cell surface CD47 by anti-human IgG Fc mAb staining (data representative of n=6 donors). (**M**) Phagocytosis of T cells coated with secreted CV1 SiRPα-Fc by NSG BMDMs in vitro (n=3). (**N**) Phagocytosis of T cells coated with secreted CV1 SiRPα-Fc by MDMs in vitro (n=7). Statistical analysis by one-way analysis of variance (ANOVA) (B, D and J), two-way ANOVA (F left, G left, I and K), Mantel-Cox (F right and G right) or unpaired, two-tailed t test (M and N) with correction for multiple comparisons by post hoc Tukey’s test (B, D and J, as well as F and G: CV1 SiRPα-Fc vs TCR + inSiRPα-Fc, I and K) or post hoc Sidak’s test (F and G: TCR + inSiRPα-Fc vs TCR + CV1 SiRPα-Fc). ****P< 0.0001; ***P < 0.001; **P < 0.01; *P < 0.05.

Finally, a side-by-side comparison of the 3 co-culture conditions, (a) tumor cells and macrophages plus recombinant CV1 SiRPα-Fc, (b) tumor cells, macrophages and CV-Fc-engineered T cells, and, (c) tumor cells and macrophages plus CV1 SiRPα-Fc supernatant (depicted in Supplemental Figure 3G), revealed that for some donors the extent of phagocytosis achieved was superior with CV1 SiRPα-Fc culture supernatants than the addition of recombinant CV1 SiRPα-Fc (provided at 10μg/ml) (Supplemental Figure 3H).

In summary, we demonstrated the secretion of SiRPα-Fc decoys by gene-modified T cells and the ability of the high-affinity CV1 SiRPα-Fc variant to augment human MDM-mediated phagocytosis of tumor cells.

### Enforced expression of CV1 SiRPα-Fc by TCR-T cells improves the control of tumor outgrowth but in a subcutaneous ACT model the T cells are depleted

Next, we performed an in vivo Winn assay in which NSG mice are subcutaneously co-injected with tumor cells plus the gene-modified T cells, and control of tumor outgrowth is evaluated over time (Figure 3E). We chose NSG mice as a model based on the high-affinity cross-species reactivity of NSG SiRPα with human CD47 (51). Consistent with our in vitro experiments, we observed significantly improved control of tumor outgrowth and survival in the context of both A375 (Figure 3F) and Me275 (Figure 3G) for CV1 SiRPα-Fc coengineered A97L-TCR T cells as compared to inSiRPα-Fc expressing ones. Complete lack of tumor control and poor survival was observed upon treatment with T cells engineered with decoys only (i.e., no tumor-specific TCR) (Figure 3, F and G).

Encouraged by these in vivo data, we subsequently sought to test the coengineered T cells in a subcutaneous tumor model (Figure 3H). Strikingly, tumor control was abrogated by TCR-T cells expressing the high affinity decoy but not the inactive one (Figure 3I). Inspection of the blood, spleen, lung, and tumors post-ACT revealed a depletion of CV1 SiRPα-Fc coengineered T cells (+/-TCR) (Figure 3J). We further observed that co-administration of recombinant CV1 SiRPα-Fc with A97L-TCR T cells resulted in complete tumor escape (Figure 3K). We reasoned that this may be due to binding of the high-affinity decoy to CD47 on the T-cell surface and subsequent macrophage-mediated depletion via the Fc tail. Indeed, flow cytometric analysis revealed T-cell surface coating by CV1-but not by inSiRPα (Figure 3L) and co-culture assays revealed phagocytosis by both NSG BMDM and human MDM of T cells coengineered to express CV1-but not inSiRPα-Fc (Figure 3, M and N).

### Neither activation-inducible expression of CV1 SiRPα-Fc nor lower affinity Fc can prevent T-cell depletion

In an effort to rescue the CV1 SiRPα-Fc coengineered T cells from depletion in vivo, we next generated lentiviral vectors encoding the decoys under the activation inducible promoter nuclear factor of activated T cells (NFAT_6_) response elements fused to the IL-2 minimal promoter (6xNFAT) to limit their expression to the TME (Figure 4A). We hypothesized that production of the decoy within the TME would favor decoy binding to tumor-cell surface CD47 and overcome peripheral depletion of the coengineered T cells. This strategy of activation-inducible gene-cargo expression has been used by others to alleviate systemic toxicity by coengineered T cells (52–54). We also optimized a co-transduction strategy for constitutive retroviral-mediated expression of the A97L-TCR and lentiviral-mediated activation-inducible upregulation of the gene-cargo in such a way that the T cells were not coated in decoy during the manufacturing process (it is critical to remove the activation beads on day 3 instead of day 5; Figure 4B) but rather upon deliberate T-cell activation (Figure 4, C-F). However, while this approach transiently improved tumor control by the CV1 SiRPα-Fc coengineered TCR-T cells (Figure 4G), they were nonetheless depleted while the non-binding inSiRPα-Fc coengineered TCR-T cells persisted (Figure 4H).

**Figure 4.**
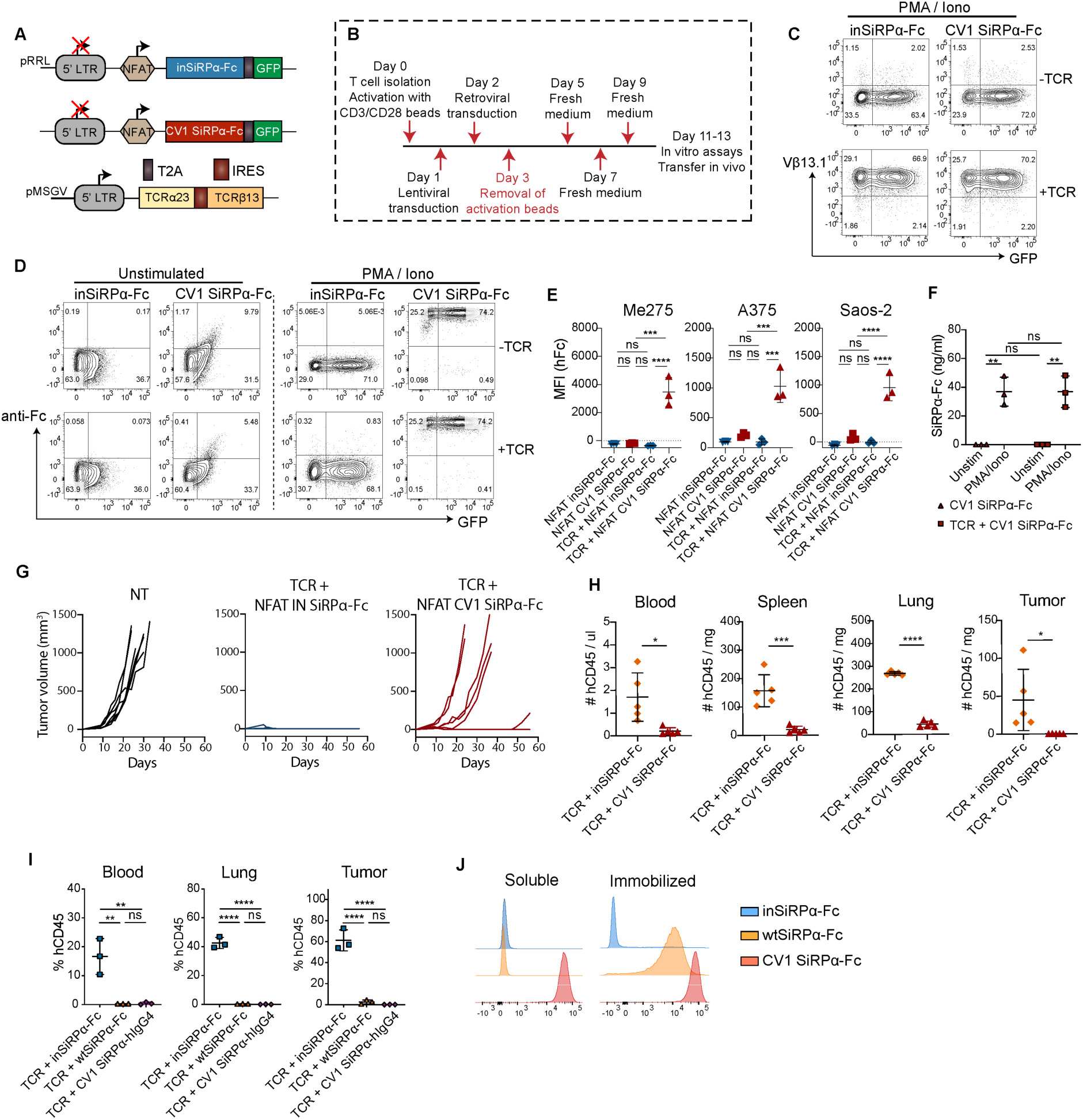
SiRPα decoys expressed under the 6xNFAT promoter are efficiently secreted upon T-cell activation. **(A)** Schematic of lentiviral constructs encoding SiRPα-Fc decoys under 6xNFAT and of a retroviral construct encoding the A97L TCR. **(B)** Strategy for optimized T cell activation, transduction, and expansion to minimize decoy secretion pre-ACT. **(C)** Detection of SiRPα-Fc and A97L-TCR by eGFP and anti-Vb13 mAb staining, respectively, in rested engineered T cells re-activated with PMA-Ionomycin for 48h (data representative of n=3 donors). **(D)** Secreted CV1 SiRPα-Fc detected on the surface of PMA-Ionomycin activated T cells versus resting T cells (data representative of n=3 donors). Gates have been placed based on non-transduced T cells. **(E)** Flow cytometric evaluation of CV1 SiRPα-Fc expression upon coculture of engineered A97L-TCR T cells with target tumor cells (n=3). **(F)** CD47-based ELISA evaluation of CV1 SiRPα-Fc secretion by engineered T cells upon PMA-Ionomycin activation (n=3). **(G)** Tumor control curves upon ACT with A97L-TCR T cells expressing SiRPα decoys under 6xNFAT (n=7). **(H)** Ex vivo evaluation of T cells persisting in the blood, spleen, lung, and tumors of NSG mice 7 days post ACT (n=5). **(I)** Frequency of human CD45^+^ cells in harvested tissues 5 days post-ACT with wtSiRPα-Fc and CV1 SiRPα-hIgG4-Fc T cells (n=3). **(J)** Binding of soluble and immobilized recombinant SiRPα-Fc οn A375 tumor cells (data representative of n=3 independent studies). Statistical analysis by one-way analysis of variance (ANOVA) (E-F and I) or unpaired, two-tailed t test (H) with correction for multiple comparisons by post hoc Tukey’s test (E-F and I). ****P< 0.0001; ***P < 0.001; **P < 0.01; *P < 0.05.

We further questioned whether substitution of human IgG1-Fc (47) with IgG4-Fc, which binds Fc receptors (FcRs) more weakly (55), on the CV1 construct could rescue the coengineered T cell from depletion, but this was not the case (Figure 4I). Moreover, TCR-T cells coengineered to express wtSiRPα-Fc decoy were also depleted in vivo (Figure 4I), despite that we were unable to detect its binding to the surface of tumor cells in vitro (Figure 2L). We hypothesized that there may be an avidity effect in vivo as multiple wtSiRPα-IgG1-Fc bound to a tumor cell are engaged by multiple FcRs on a macrophage. Indeed, we showed that immobilization of wtSiRPα-Fc on protein G-coated beads enabled tumor-cell binding levels similar to the high-affinity decoy (Figure 4J).

Thus, in summary we demonstrated that T cells can be coengineered to express both a tumor-redirected TCR and CV1 SiRPα-Fc decoys that potentiate macrophage-mediated phagocytosis of tumor cells in vitro. Moreover, in a Winn assay, such co-engineered T cells significantly improve the control of tumor outgrowth and mouse survival. However, CV1 SiRPα-Fc coats the surface of T cells themselves leading to their depletion upon ACT in subcutaneous tumor models, regardless of whether the decoy is constitutively or inducibly expressed under 6xNFAT. T cells coengineered to express wtSiRPα-Fc or CV1 SiRPα-IgG4-Fc are also depleted in vivo. Hence, the gene-modification of T cells with SiRPα-Fc decoys (or likely any anti-CD47 molecule fused to an Fc) is not a viable option for clinical translation.

### Tumor-specific antibodies synergize with CV1 SiRPα monomers to potentiate macrophage-mediated phagocytosis of tumor cells

Previous studies have demonstrated that CD47 blockade with monomeric SiRPα decoys can lower the phagocytic threshold, but macrophage mobilization and target cell engulfment further requires pro-phagocytic signals such as via Fc/FcR engagement (33, 38). We thus reasoned that we could remove the prophagocytic IgG1 Fc tail from the CV SiRPα decoy to spare the T cells from depletion and instead co-administer a tumor targeted monoclonal antibody (mAb) comprising an active Fc tail (Figure 5A).

**Figure 5.**
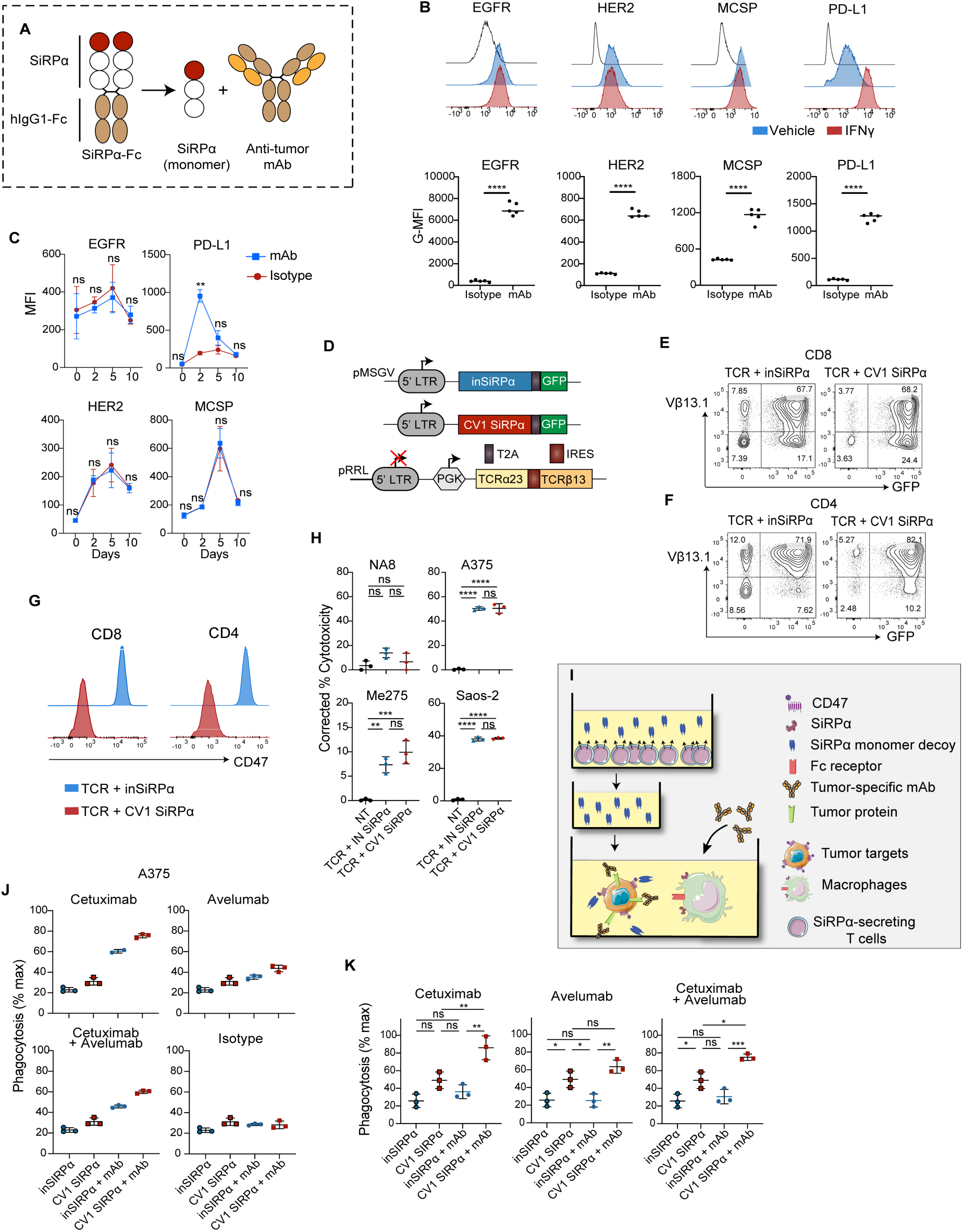
Tumor-targeted monoclonal antibodies synergize with T-cell secreted CV1 SiRPα monomer decoys to augment tumor cell phagocytosis by macrophages in vitro. **(A)** Schematic of strategy to combine SiRPα monomer with tumor-targeted mAbs. **(B)** Expression of EGFR, HER2, MCSP and PD-L1 on the surface of A375 tumor cells in vitro (top) and in established tumors in vivo (bottom) (n=5). **(C)** Evaluation of EGFR, HER2, MCSP and PD-L1 expression by cultured CD8^+^ T cells at different timepoints (n=3). **(D)** Schematic of retroviral vectors encoding SiRPα monomers and of lentiviral vector encoding the A97L TCR. **(E)** Expression of SiRPα monomer and A97L-TCR in transduced CD8^+^ and, **(F)** CD4^+^ T cells, detected by eGFP and anti-Vβ13.1 mAb staining, respectively (data representative of n=11 donors). **(G)** Flow cytometric detection of CV1 SiRPα monomer binding on CD8^+^ and CD4^+^ T cells by comparison of anti-CD47 mAb staining to CV1-versus inSiRPα-engineered T cells (data representative of n=11 donors). **(H)** Frequency of Annexin V^+^ DAPI^+^ cells tumor cells in 24h co-cultures with SiRPα monomer coengineered A97L-TCR T cells at E:T = 1:1, corrected to tumor alone (n=3). **(I)** Schematic of macrophage-mediated tumor cell phagocytosis assay in the presence of SiRPα monomer and Cetuximab or/and Avelumab. **(J)** Human MDM phagocytosis of A375 tumor cells in the presence of T-cell secreted SiRPα monomer alone versus in combination with Cetuximab or/and Avelumab (representative results from n≥3 donors). **(K)** NSG murine BMDM phagocytosis of A375 tumor cells in the presence of T cell-secreted SiRPα monomer alone versus in combination with Cetuximab or/and Avelumab (n=3). Statistical analysis by unpaired, two-tailed t test (B, bottom panel), two-way analysis of variance (ANOVA) (C) or one-way ANOVA (H and K) with correction for multiple comparisons by post hoc Sidak’s test (C) or post hoc Tukey’s test (H and K). ****P< 0.0001; ***P < 0.001; **P < 0.01; *P < 0.05.

We identified epidermal growth factor receptor (EGFR), human epidermal growth factor receptor 2 (HER2), melanoma-associated chondroitin sulfate proteoglycan (MCSP) and programmed death ligand 1 (PD-L1) as being homogeneously expressed by A375 tumor cells in vitro (Figure 5B upper panel) and in vivo in established tumors (Figure 5B lower panel). EGFR, HER2 and MCSP are all cell-surface receptors commonly deregulated in cancer and PD-L1 is a checkpoint receptor frequently upregulated by tumors (56). We acquired the clinically available mAbs Cetuximab, Trastuzumab and Avelumab, targeting EGFR, HER2 and PD-L1, respectively, which each comprise a functional IgG1 Fc tail needed for antibody dependent cellular phagocytosis (ADCP) by macrophages. To target MCSP, an in-house human IgG1 Fc fusion antibody was generated using publicly available scFv sequences. Aside from transient upregulation of PD-L1 during activation (57), none of the antigens were present on the surface of cultured CD8^+^ and CD4^+^ T cells (Figure 5C, Supplemental Figure 4A).

We next built retroviral vectors encoding the SiRPα monomers and GFP (Figure 5D). Using our optimized protocol (Figure 2B), we achieved high co-transduction efficiencies of both CD8^+^ (Figure 5E) and CD4^+^ (Figure 5F) T cells as measured by GFP expression and anti-Vβ13.1 Ab staining. Lower anti-CD47 antibody cell-surface staining was observed for CV1 versus inSiRPα coengineered TCR T cells indicative of CD47 masking by binding of the high affinity decoy (Figure 5G). As previously observed for CV1 SiRPα-Fc, constitutive secretion of CV1 SiRPα monomers did not affect the cytotoxic capacity of the A97L-TCR T cells (Figure 5H).

Subsequently, we set up a series of co-culture assays comprising macrophages and tumor cells along with CV1 or control inSiRPα decoys (collected from culture supernatants of engineered T cells) and the different tumor-specific mAbs (Figure 5I, Supplemental Figure 4B). We observed significant increases in A375 phagocytosis by MDMs in the presence of Cetuximab or/and Avelumab along with CV1 SiRPα monomer, as compared to phagocytosis in the presence of CV1 alone (Figure 5J). Anti-MCSP mAb whereas did not synergize with CV1, and Trastuzumab did not facilitate ADCP of A375 (Supplemental Figure 4C). The same trends were observed for co-culture assays comprising Saos-2 (Supplemental Figure 4D). A non-tumor cell binding isotype control mAb had no impact on phagocytosis (Figure 5J bottom right). Moreover, in co-culture assays of either A375 (Figure 5K) or Saos-2 (Supplemental Figure 4E) tumor cells with bone marrow derived macrophages (BMDMs) generated from NSG mice, CV1 SiRPα monomer potentiated phagocytosis and synergized with Cetuximab or/and Avelumab (Figure 5K, Supplemental Figure 4E).

Finally, taking into consideration that Avelumab is a checkpoint inhibitor we sought to test if PD-L1 blockade had any impact on the effector function of T cells in co-culture with tumor cells. Under the in vitro conditions tested, neither target cell killing, nor IFNγ production by T cells were altered in the presence of Avelumab (or Cetuximab; Supplemental Figure 4, F and G).

In summary, we demonstrated efficient coengineering of A97L-TCR T cells with monomeric SiRPα decoys and we showed that both Cetuximab and Avelumab synergize with them to augment tumor-cell mediated phagocytosis by human MDMs.

### Human T cells coengineered to inducibly secrete SiRPα monomers are depleted in vivo

Encouraged by our findings, we next built lentiviral vectors encoding CV1- and inSiRPα monomer, each along with GFP as a marker of expression, under 6xNFAT (Figure 6A). We efficiently co-transduced T cells to express the A97L-TCR and the inducible decoys and demonstrated expression upon T-cell co-culture with target cells (Figure 6B). However, we observed superior A375 tumor control by TCR-T cells expressing the inactive decoy (Figure 6C), indicative of T-cell depletion upon expression of CV1 SiRPα monomer. Others have recently coengineered chimeric antigen receptor (CAR)-T cells to secrete a truncated monomeric CV1 SiRPα ectodomain (encompassing residues 27-118 versus 27-371) and reported T-cell persistence (58). We, however, observed depletion of T cells engineered to express either of these CV1 molecules, albeit at a slower rate over days for ones expressing the truncated version (Supplemental Figure 5A).

**Figure 6.**
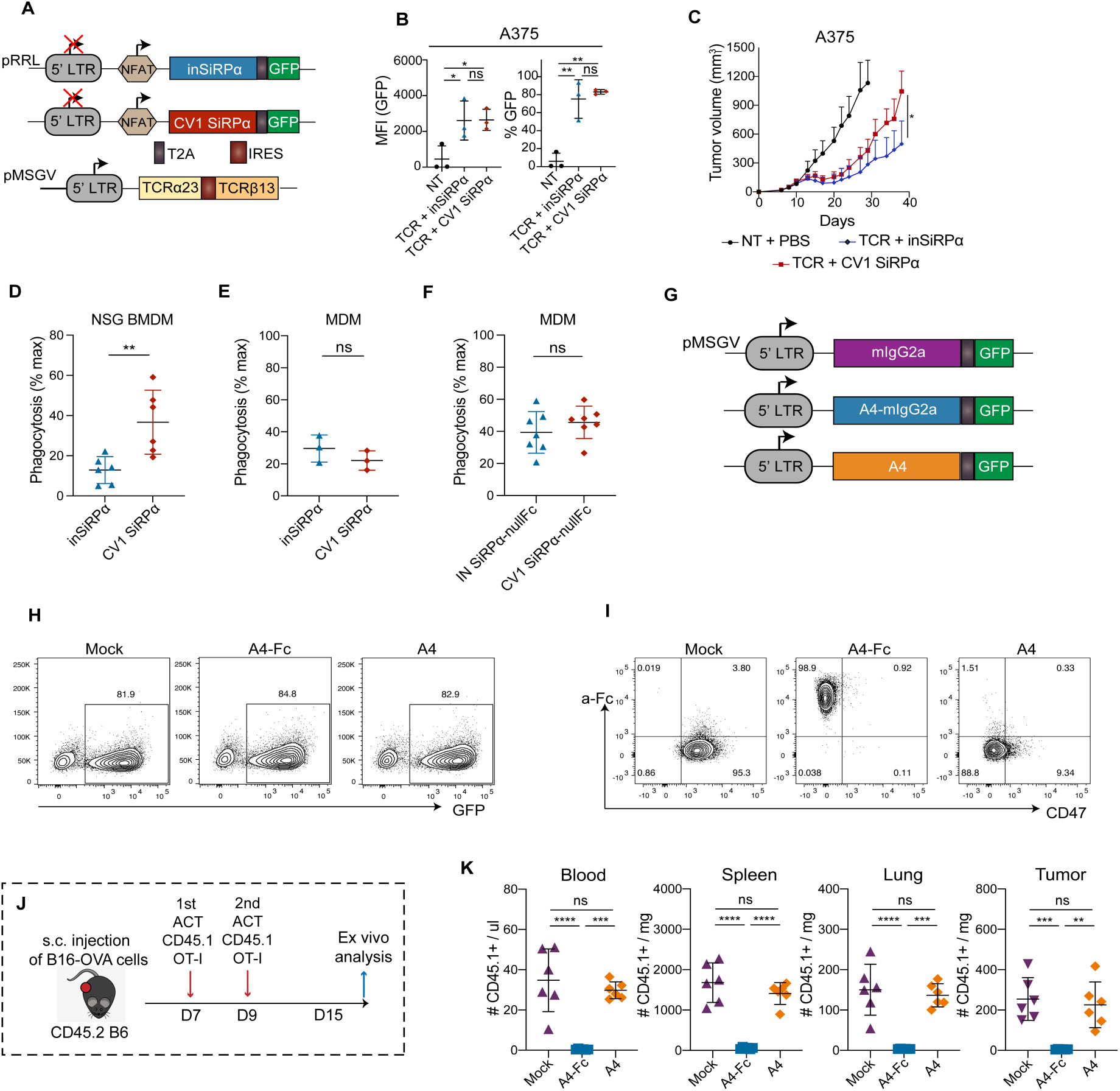
Monomeric SiRPα decoy-coated human T cells are targeted for phagocytosis by macrophages of NSG but not of human origin, and murine T cells are only depleted C57BL/6 mice if the CD47 decoy is fused to an active Fc tail. (**A**) Schematic of lentiviral vectors encoding SiRPα monomer under 6xNFAT and of a retroviral vector encoding the A97L-TCR. (**B**) Inducible SiRPα monomer expression by A97L-TCR T cells as detected by eGFP expression upon co-culture with target tumor cells (n=3). **(C)** A375 tumor control curves following ACT with A97L-TCR T cells coengineered to express inSiRPα or CV1 SiRPα monomers under 6xNFAT (n≥6, data representative of 2 independent studies). **(D)** Evaluation of phagocytosis of A97L-TCR T cells coengineered to express inSiRPα or CV1 SiRPα monomers by BMDMs (n=6). **(E)** Evaluation of phagocytosis of A97L-TCR T cells coated with inSiRPα or CV1 SiRPα monomers by human MDMs (n=3). **(F)** Evaluation of phagocytosis of T cells coated with secreted CV1 SiRPα-null-Fc by human MDMs in vitro (n=7). **(G)** Schematic of retroviral constructs encoding A4-Fc and A4 decoys. **(H)** Expression of A4-Fc and A4 decoys in transduced mouse OT-I T cells, detected by eGFP (data representative of n≥3 donors). **(I)** Flow cytometric detection of A4-Fc and A4 monomer binding on OT-I T cells by staining with anti-Fc mAb and anti-mouse CD47 mAbs (data representative of n=3 donors). **(J)** Schematic of ACT against subcutaneous B16-OVA tumors and ex vivo analysis. **(K)** Frequency and number of mouse CD45.1^+^ cells in harvested tissues 6 days post-ACT (n=6, data representative of 2 independent studies). Statistical analysis by one-way analysis of variance (ANOVA) (B and K), two-way ANOVA (C), or unpaired, two-tailed t test (D-F) with correction for multiple comparisons by post hoc Tukey’s test (B and K) or post hoc Sidak’s test (C). ****P< 0.0001; ***P < 0.001; **P < 0.01; *P < 0.05.

In subsequent co-culture assays we observed that T cells coengineered to express CV1 SiRPα (aa 27-371) monomer are phagocytosed by macrophages of NSG origin (BMDM, Figure 6D) but, importantly, not by ones of human origin (MDM, Figure 6E). We further demonstrated that human T cells engineered to express CV1 SiRPα fused to an inactive Fc tail (null) are not phagocytosed by MDMs (Figure 6F). Thus, our data suggest that there is potential for the clinical translation of human T cells coengineered to express molecules that block CD47, provided that they do not comprise an active Fc tail.

We hypothesized that human MDMs do not phagocytose T cells coengineered to express CV1 SiRPα monomers (or CV1 SiRPα-nullFc) because, despite that CD47 is blocked, other ‘don’t eat me’ signals are at play such as CD24 (59), β2m (60) and PD-L1(61), thus elevating the phagocytic threshold. We further speculated that the persistence of human T cells in NSG mice is critically dependent upon human CD47 receptor engagement by SiRPα receptor on the surface of NSG myeloid cells such that mere blockade of the axis drives macrophage-mediated phagocytosis. Indeed, the murine macrophage receptor LILRB1 does not cross-react with human β2m (60) and we observed no PD-1 on tumor-infiltrating macrophages or neutrophils pre-ACT (Supplemental Figure 5B). Thus, 2 major ‘don’t eat me’ axes are absent in the xenograft model. We did, however, detect Siglec-G (the murine homologue of Siglec-10) on the surface of tumor infiltrating macrophages (Supplemental Figure 5C), which, along with its ligand CD24, constitutes a recently identified ‘don’t eat me’ axis (59). We questioned whether the overexpression of CD24 on human T cells could rescue them from depletion. To that end, we built retroviral vectors encoding murine or human CD24 (we tested both due to low sequence homology) and the A97L TCR (Supplemental Figure 5D). We coengineered human T cells with the A97L-TCR and CD24 along with CV1- or inSiRPα monomer and tested them in vivo. The overexpression of CD24 did not circumvent SiRPα-mediated depletion of T cells (Supplemental Figure 5E&F) and compromised tumor control by TCR-T cells (Supplemental Figure 5G). We did not explore this further.

Finally, we sought to test the impact of engineering murine T cells with a soluble decoy of SiRPα versus one fused to an active Fc tail, predicting that the latter would lead to T-cell depletion in C57BL/6 mice. We built retroviral constructs encoding a previously described nanobody A4 targeting CD47 and fused or not to mIgG2a (62). We did not use CV1 as it has been reported to have weaker binding affinity for murine than human CD47 (Figure 5G) (62). Murine OT1 TCR T cells were efficiently transduced with the different constructs (Figure 5H) and expression detected by flow cytometry (anti-Fc staining or decrease in anti-CD47 staining, Figure 5I). The engineered T cells were adoptively transferred into C57BL/6 mice bearing subcutaneous B16-OVA tumors. While the A4-Fc T cells were depleted in the blood, spleen, lung and tumor, the mock and A4-engineered T cells persisted. These data further support our assertion that there is potential for the clinical translation of tumor-redirected T cells expressing monomeric decoys of CD47.

### Cetuximab and Avelumab synergize with TCR-T cells to reprogram the TME and improve tumor control

While generating the viral constructs and performing the in vivo studies described above for the ACT of human TCR-T cells coengineered to inducibly secrete CV1 SiRPα monomer in tumor-bearing NSG mice, we tested Cetuximab or/and Avelumab (both comprising active human IgG1 Fc tails) (56) in vivo. We began with Winn assays (Figure 7A) and observed that the mAbs on their own did not slow tumor outgrowth (Figure 7B). However, the coadministration of Cetuximab or/and Avelumab with TCR-T cells further delayed tumor growth mediated by TCR-T cells alone (Figure 7C). In addition, co-administration of the mAbs with TCR-T cells led to significantly prolonged survival of the mice (Figure 7D). Finally, in a subcutaneous tumor model (Figure 7E) we observed that administration of the mAbs (both individually and in combination) had no impact on the growth of established tumors (Figure 7F). However, the mAbs individually and together enabled significantly improved tumor control in combination with ACT (Figure 7G). Interestingly, an evaluation of subcutaneous tumor control by A97L-TCR T cells in the presence of full IgG versus F(ab’)2 fragments of Cetuximab or Avelumab revealed a role for the Fc tail in tumor control when targeting EGFR but not PD-L1 (Supplemental Figure 6, A and B).

**Figure 7.**
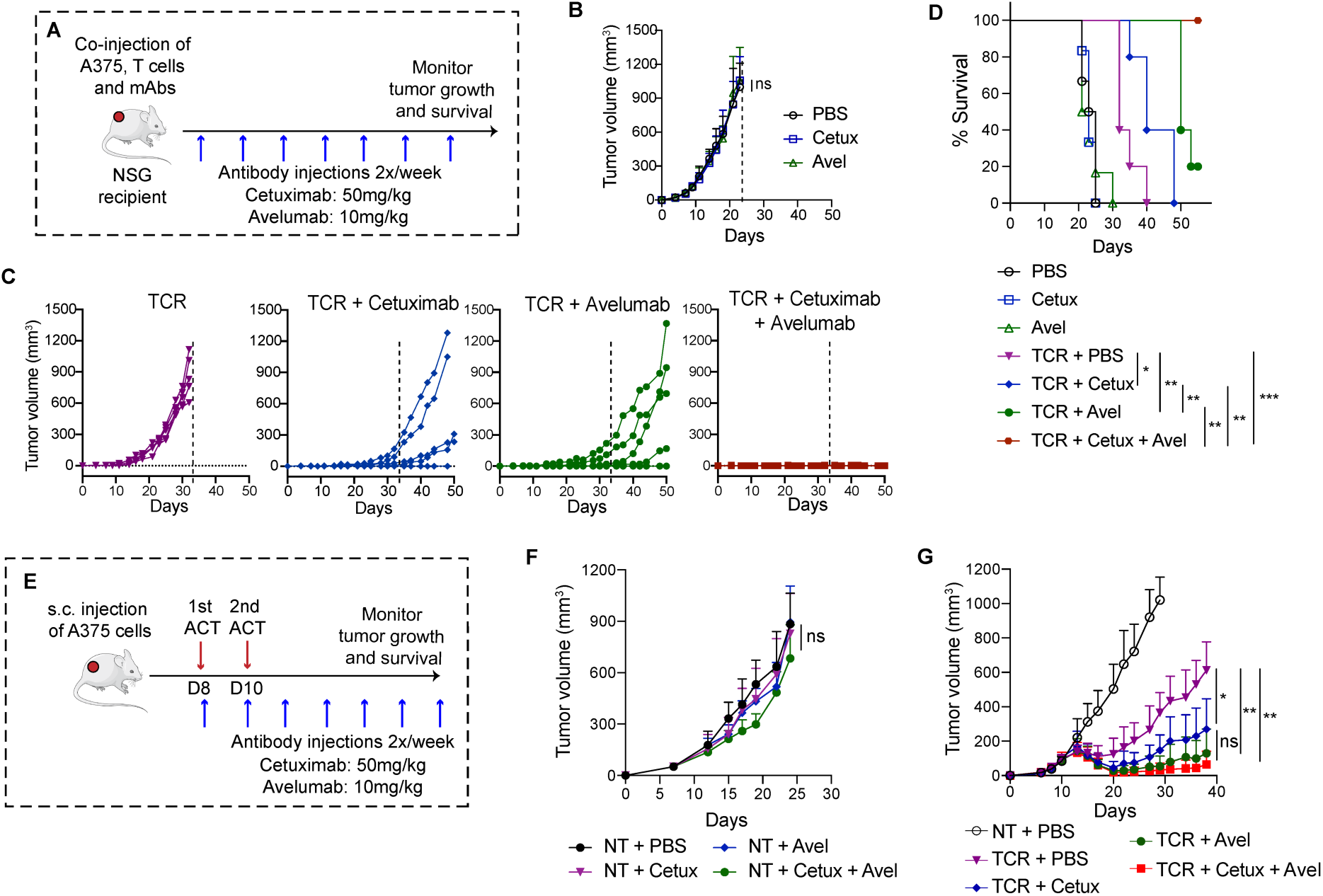
Enhanced tumor control upon coadministration of tumor-targeted monoclonal antibodies with A97L-TCR T cells. **(A)** Schematic of Winn assay for A375 tumor cells with A97L-TCR T cells co-administered with Cetuximab or/and Avelumab targeting EGFR and PD-L1, respectively. **(B)** Winn assay with tumor-targeted mAbs alone, with **(C)** A97L-TCR T cells alone, or for A97L-TCR T cells in combination with Cetuximab, or/and Avelumab (n≥5, data representative of 2 independent studies). **(D)** Survival curves for the Winn assay. **(E)** Schematic of ACT and tumor-targeted mAb co-administration against subcutaneous Saos-2 tumors. **(F)** Treatment of established A375 tumors with tumor-targeted mAbs alone (n=7, data representative of 2 independent studies). **(G)** A375 tumor control following ACT with A97L-TCR T cells alone versus in combination with Cetuximab or/and Avelumab (n≥5, data representative of 3 independent studies). Statistical analysis by two-way analysis of variance (ANOVA) (B, F and G) or Mantel-Cox (D) with correction for multiple comparisons by post hoc Tukey’s test (B, F and G). ****P< 0.0001; ***P < 0.001; **P < 0.01; *P < 0.05.

Ex vivo characterization of tumors from the differently treated mice (Supplemental Figure 6C) revealed a significant increase in the infiltration of adoptively transferred CD8^+^ and CD4^+^ T cells (Supplemental Figure 6D) upon coadministration of both Cetuximab and Avelumab, as well as a higher CD8:CD4 T cell ratio (Supplemental Figure 6E). Notably, the coadministration of Avelumab (alone or with Cetuximab) was associated with higher proliferative capacity of T cells as measured by Ki-67 staining (both frequency and MFI, Supplemental Figure 6, F and G). We also observed a higher frequency of central memory CD8^+^ and CD4^+^ T cells (T_CM_) (Supplemental Figure 6H), and significantly lower expression (both percentage and MFI) of the inhibitory markers PD-L1, TIM-3 and PD-1 on CD8^+^ and CD4^+^ T cells, from the tumors of mice treated with both mAbs as compared to ACT alone (Supplemental Figure 6, I and J).

We also observed changes to the endogenous myeloid compartment upon coadministration of Avelumab and Cetuximab (Supplemental Figure 7A). For example, we detected higher intratumoral presence of CD11c^+^ DCs, F4-80^+^ macrophages, and F4-80^int^ Ly6C^+^ monocytic-derived macrophages, but reduced numbers of Ly6G^+^ neutrophils, both in terms of absolute numbers (Supplemental Figure 7B) but also proportionally within the mouse myeloid compartment in favor of macrophages (Supplemental Figure 7C). Increased macrophage numbers were accompanied by phenotypic changes, including exhibiting a more M1-like phenotype as evaluated by staining for CD38 and Egr2 (Supplemental Figure 7D).

Although it was not possible to test the combination of affinity-optimized TCR-T cells coengineered to express CV1 SiRPα monomer along with Cetuximab and Avelumab in a pre-clinical tumor model, taken together we have presented strong evidence that our strategy will be feasible and effective in humans. Indeed, we demonstrated in vitro that human MDMs do not phagocytose human T cells expressing CV1 SiRPα monomer and that Cetuximab and Avelumab synergize with the decoy to augment target cell phagocytosis. In addition, Cetuximab and Avelumab in combination with ACT favorably support T cells and harness endogenous innate immunity, including higher intratumoral presence of M1-like macrophages which would presumably be responsive to CD47 blockade.

## DISCUSSION

Although the primary target of immunotherapy is arguably adaptive immunity, in particular cytotoxic T lymphocytes having the capacity to directly kill tumor cells, the development of approaches for harnessing innate immunity (19, 21, 63, 64) is gaining momentum and is imperative for improving patient responses. In addition, strategies for circumventing potential toxicity, which will likely only increase in the context of combination strategies, are paramount for safely advancing efficacious therapies to the clinic (65, 66). Here, with the aim of improving the control of solid tumors, we have developed a novel combinatorial ACT strategy that directly optimizes the activity of TCR-engineered T cells and exploits the phagocytic capacity of endogenous macrophages.

For our study, we sought to utilize affinity-optimized TCR-engineered T cells as a vehicle for delivering SiRPα decoys to block CD47 directly in the TME. Indeed, important clinical failures, largely due to severe hemolytic reactions caused by hemagglutination induced by anti-CD47 mAb (i.e., systemic administration), have been reported (67). We began by gene-engineering CD8^+^ and CD4^+^ T cells with an A2/NY-specific TCR comprising a single amino acid replacement (A97L) which increases its binding strength to within the upper range of natural TCR affinity and augments in vitro effector functions as compared to WT (9, 41). Our findings are in line with numerous studies demonstrating that TCR strength is a key determinant of T cell response and that TCRs of ‘intermediate affinity’ confer maximal effector function (8, 68–72). Here, we have demonstrated higher proliferative capacity of the affinity-optimized A97L-TCR T cells in the TME following ACT, as well as significantly improved solid tumor control and survival. In future studies, we seek to elucidate the molecular mechanisms behind the improved functionality of affinity-optimized A97L-versus WT-TCR engineered T cells.

Macrophages play an important role in the removal of tumor cells from the body via phagocytosis but in most cancers high macrophage infiltration is associated with poor survival (73). Moreover, macrophages can reversibly alter their endotype in response to environmental cues including from parenchymal and immune cells and the extracellular matrix. Hence, macrophages represent a promising target in cancer immunotherapy. Here, we have focused on the CD47/SiRPα ‘don’t eat me’ axis as it has been shown to constitute a powerful immune checkpoint of macrophages in the context of both liquid and solid tumors (74). While not addressed in our study, our approach may further benefit by combination with treatments that can modulate the myeloid compartment such as low-dose radiotherapy (21) or colony stimulating factor 1 receptor (CSF1R) inhibitors (75) that can drive tumor infiltration or/and polarize macrophages from an anti-inflammatory M2-towards a tumoricidal M1-endotype (76).

Efficient gene-modification of human T cells with both the A97L-TCR and the different SiRPα decoys was achieved by utilizing a dual lentivirus and retrovirus transduction strategy, and we demonstrated that CV1 SiRPα-Fc decoy secreted by T cells could significantly improve macrophage mediated phagocytosis of tumor cells. However, we observed that the engineered T cells themselves were coated in the high-affinity decoy and were susceptible to phagocytosis by both human MDMs and mouse BMDMs in vitro. Moreover, T cells coengineered to secrete CV1 SiRPα-Fc decoy were depleted in vivo. Importantly, in terms of potential for clinical translation, CV1 SiRPα monomer (or comprising a null Fc tail) engineered T cells were spared from phagocytosis by human MDMs in vitro, presumably due to the presence and recognition of other ‘don’t eat me’ signals on the T cells.

It has been well established that although CD47 blockade can lower the phagocytic threshold, prophagocytic signals such as via Fc/FcR engagement are needed to drive target cell engulfment by macrophages (33, 38). We identified the clinically approved mAbs Avelumab and Cetuximab, both comprising active Fc tails, as capable of binding specifically to the target cell line A375 and of synergizing with CV1 decoys to augment tumor cell phagocytosis in vitro. Both mAbs significantly improved subcutaneous A375 tumor control upon ACT with A97L-TCR T cells. Interestingly, tumor control upon ACT was negatively impacted by removal of the Fc tail from Cetuximab but not from Avelumab. Indeed, Avelumab is cross-reactive with murine PD-L1 which is frequently upregulated on stromal and immune cells in the TME (77) and hence may contribute to tumor control in our studies via immune checkpoint blockade. Notably, a growing body of clinical data supports the combinatorial use of Cetuximab and Avelumab (78). For example, the treatment of metastatic colorectal cancer patients with Cetuximab drives CD8^+^ T cell infiltration and the upregulation of PD-L1, the latter of which can be countered with Avelumab (79) to support the infiltrating T cells. Similar observations have been made in pre-clinical studies (80).

Due to the apparent nonredundant role played by the human CD47/ murine SiRPα axis in enabling human T cell engraftment in NSG mice (51), it was not possible for us to evaluate our coengineering strategy in the context of subcutaneous tumor models because the T cells were blocked by CV1 and depleted in vivo, even if the decoys were inducibly expressed as a monomer under 6xNFAT. In Winn assays, however, in line with our in vitro co-culture data, we observed significantly improved control of tumor outgrowth and survival of mice treated with CV1 SiRPα-Fc coengineered A97L-TCR T cells versus A97L-TCR T cells. Notably, Dacek *et. al*. recently reported that CD19 CAR-T cell transfer augmented anti-CD20 Rituximab therapy through innate immune activation and that responses could be improved upon coengineering with CV1 (58). In contrast to our in vivo experiments, they observed persistence of their gene-modified T cells. One important difference between our studies is the use of a TCR versus CAR, but others recently revealed that CD47 expression is critical for CAR T-cell survival in vivo (81). Also of mention, in another study murine CAR-T cells coengineered to secrete murine wtSiRPα-Fc (derived from mouse IgG2a) were reported to augment tumor control (82). We, on the other hand, observed depletion of murine OT1 TCR T cells coengineered to secrete the decoy A4 fused to the same Fc (but not for A4 alone). Possibly, SiRPα decoy engineered T cells may persist in vivo in one study but not in another due to due to higher proliferative capacity of the T cells, or/and due to differences in the phagocytic threshold of the endogenous murine macrophages.

Taken together, we have shown that human T cells can be effectively engineered to secrete functional high-affinity SiRPα decoys of CD47. Moreover, our in vitro data indicate that T cells coengineered to secrete high-affinity decoy monomer (i.e., no active Fc tail) would not be depleted upon ACT in humans, presumably due to the presence of other ‘don’t eat me’ signals (59). In addition, we have demonstrated that coadministration of Avelumab and Cetuximab enhances tumor control upon ACT and is associated with favorable changes to innate immunity including of macrophages which may be responsive to local blockade of the SiRPα/CD47 axis. Without question, it will take careful consideration and dose escalation studies to safely deliver combinations of ACT with checkpoint blockade of both adaptive and innate immunity, but the strategy holds considerable potential for tackling difficult to treat solid and liquid tumors alike.

## MATERIALS AND METHODS

### Study design

The objective of this study was to develop a combination ACT therapy comprising T cells bearing an affinity-optimized TCR targeting A2/NY and coengineered to secrete high-affinity SiRPα binding domain decoys (CV1) in order to harness macrophages (i.e., via CD47 blockade) in the TME to augment tumor control. T-cell transduction was achieved by both lentiviral and retroviral vectors. The decoys were first engineered in T cells as Fc fusion proteins, then as Fc fusion proteins but expressed under 6xNFAT to allow activation inducible secretion, and finally as monomers and combined with tumor-targeting mAbs. The binding properties and functional capacity of recombinant CV1 and control decoys (WT and inactive) were validated at the onset of the study by flow cytometry and co-culture measurements of phagocytosis, respectively. Experiments in NSG mice were performed with 5-9 mice per group as indicated in the figure legends on the basis of previous experiments showing that this size could guarantee good reproducibility and statistically significant differences. Tumor-bearing mice were randomized into treatment groups before T cell infusion based on the size of the established tumors. Mice were treated by an operator who was blinded to treatment groups. All in vitro experiments were performed with T cells from a minimum of 3 independent healthy donors. The number of repetitions is indicated in the figure legends. All analysis of in vitro and in vivo data was based on objectively measurable data.

### Mice

NOD.Cg-*Prkdc^scid^ Il2rg^tm1Wjl^*/SzJ (NSG) mice were obtained from the Jackson Laboratory and subsequently maintained and bred in-house under specific opportunist pathogen free (SOPF) conditions at the Epalinges UNIL animal facility. CD45.1^+^ C57BL/6 female mice were also bred in-house, while female CD45.2^+^ C57BL/6 mice aged 6-12 weeks old were purchased from Harlan (Harlan, Netherlands).

### Cell lines

The cell lines A375 (melanoma; HLA-A*0201^+^, NY-ESO-1^+^), NA8 (melanoma; HLA-A*0201^+^, NY-ESO-1^-^), Saos-2 (osteosarcoma; HLA-A*0201^+^, NY-ESO-1^+^), Jurkat (T cell leukemia), HEK293T (human embryonic kidney) and Phoenix-ECO cells were purchased from the ATCC. Me275 (melanoma; HLA-A*0201^+^, NY-ESO-1^+^) was kindly provided by Prof. Daniel Speiser (University of Lausanne). CD47-deficient Jurkat variant JinB8 cells, originally described by Dr. Eric Brown, were kindly provided by Dr. David Roberts (Center for Cancer Research, National Cancer Institute) (83). The B16-OVA melanoma tumor cell line was a kind gift from Prof. Pedro Romero (UNIL). NA8, Me275, A375 and Saos-2 cell lines were engineered with NucLight lentivirus (Incucyte) to stably express nuclear-restricted mKate2 fluorescent protein in order to track their activity in vitro, according to the manufacturer’s instructions. All human melanoma cell lines were maintained in IMDM (Thermo Fisher Scientific) supplemented with 10% FCS and 1% Penicillin/Streptomycin. All remaining cell lines were cultured in RPMI + Glutamax (Thermo Fisher Scientific) supplemented with 10% FCS and 1% Penicillin/Streptomycin. All cell lines tested negative for mycoplasma contamination.

### Molecular cloning

cDNA sequences of human wild-type and A97L TCR AV23.1 and BV13.1 genes specific for NY-ESO-1_157-165_ epitope presented by HLA-A*0201 are from (9, 42). cDNA sequences of TCR AV12 and BV13.3 genes specific for human Melan-A/MART-1_26-35_ epitope presented by HLA-A*0201 are from (45). The sequences were codon-optimized and synthesized by GeneArt (Thermo Fisher) in a single cassette separated by an internal <u>r</u>ibosomal <u>e</u>ntry <u>s</u>ite (IRES) or a 2A sequence self-cleaving peptide sequence. Subsequently, the cassettes were cloned into pRRL lentiviral and pMSGV retroviral vectors using standard molecular cloning techniques. To create SiRPα-Fc fusion soluble proteins the cDNA sequence of the full extracellular domain (aa 27-371) of human wild-type allele 1 SiRPα (wtSiRPα) cDNA (Uniprot: P78324) was codon-optimized and synthesized by GeneArt (Thermo Fisher Scientific), fused with the cDNA sequence of the human hinge and IgG1 heavy chain Fc region (pFUSE-hIgG1-Fc1, InvivoGen), downstream of a mouse Ig kappa chain secretion signal peptide. CV1 high-affinity SiRPα (CV1 SiRPα) sequence is from (38). A non-binding SiRPα (inSiRPα) variant sequence is from (48). The cassettes were subsequently cloned into a pXLG expression vector under the control of a CMV promoter, kindly provided by Prof. David Hacker at the Technological Platform for Protein Production (École Polytechnique Fédérale de Lausanne). Proteins were produced in CHO cells and purified by affinity chromatography at the same platform.

For constitutive expression of decoys by T cells, the sequences of Fc-fused SiRPα molecules were subcloned into a pMSGV retroviral vector followed by enhanced (e)GFP reporter gene cassette using PCR and standard molecular cloning techniques to generate pMSGV-wtSiRPα-Fc-T2A-eGFP, pMSGV-inSiRPα-Fc-T2A-eGFP and pMSGV-CV1 SiRPα-Fc-T2A-eGFP retroviral plasmids. SiRPα monomers were generated by removing the IgG1 Fc from the aforementioned retroviral plasmids using standard molecular cloning techniques to generate pMSGV-wtSiRPα-T2A-eGFP, pMSGV-inSiRPα-T2A-eGFP, and pMSGV-CV1 SiRPα-T2A-eGFP retroviral plasmids. For inducible expression in T cells the sequences of all monomer and Fc-fused SiRPα molecules were subcloned into a pRRL lentiviral vector under the control of an inducible promoter containing 6 repeats of the NFAT response element and a minimal IL-2 promoter, as previously described (84), to generate pRRL-NFAT-inSiRPα-T2A-eGFP, pRRL-NFAT-CV1 SiRPα-T2A-eGFP, pRRL-NFAT-inSiRPα-Fc-T2A-eGFP and pRRL-NFAT-CV1 SiRPα-Fc-T2A-eGFP lentiviral plasmids.

For constitutive secretion of anti-CD47 A4-Fc fusion proteins by murine T cells the cDNA sequence of the high-affinity anti-mouse CD47 nanobody A4 (35) was codon-optimized and synthesized by GeneArt (Thermo Fisher Scientific), fused with the cDNA sequence of the mouse hinge and IgG2a heavy chain Fc region (pFUSE-mIgG2a-Fc1, InvivoGen), downstream of a mouse Ig kappa chain secretion signal peptide and was subcloned into a pMSGV retroviral vector followed by enhanced (e)GFP reporter gene cassette using PCR and standard molecular cloning techniques to generate pMSGV-A4-Fc-T2A-eGFP. A4 monomers were generated by removing the IgG2a Fc from the aforementioned retroviral plasmids using standard molecular cloning techniques to generate pMSGV-A4-T2A-eGFP.

To generate an anti-human MCSP/CSPG4 hIgG1-Fc fusion antibody-like protein, the sequence of the light chain of the anti-human MCSP/CSPG4 scFv (clone 9.2.27) was subcloned into pXLG expression vector followed by a GS linker and the sequence of the heavy chain. The cassette was placed upstream of the cDNA sequence of the human hinge and IgG1 heavy chain Fc region to generate an IgG1 Fc-fusion protein.

### Flow cytometry

For flow cytometric analysis, single cell suspensions were stained with antibodies against human CD8 (SK1, RPA-T8), CD4 (SK3, OKT4, RPA-T4), CD45RA (2H4), CD197/CCR7 (G043H7), CD69 (FN50), CD137 (4B4-1), LAG3 (3DS223H), TIM3 (F38-2E2), PD-1 (NAT105), CD47 (CC2C6), CD45 (H130), EGFR (AY13), HER2 (24D2), MCSP (EP-1), and PD-L1 (29E.2A3), and against mouse CD45(30F11), CD11c (N418), Ly6G (1A8), Ly6C (H414), F4-80 (ΒΜ8), Egr2 (erongr2), CD38 (90) and SiRPα (P84). Antibodies were purchased from BD Biosciences, Biolegend, Thermo Fisher Scientific, Miltenyi Biotec, ImmunoTools, Beckman Coulter, Sino Biological or produced in-house from hybridomas at the flow cytometry platform. Matched isotypes were used as indicated.

### Detection of CV1 SiRPα-Fc and A4-Fc binding to cell surface expressed CD47

Detection of soluble SiRPα-Fc binding on tumor cell surface CD47 was achieved by incubation of the cells with soluble human SiRPα-Fc fusion proteins (produced as described above) at 4°C for 1h, followed by staining with labeled goat anti-human IgG Fc (HP6017, Biolegend) at 4°C for 30min. To enhance the strength and stability of soluble SiRPα-Fc binding on cell surface CD47, soluble SiRPα-Fc at saturating concentrations were immobilized on 0.4-0.6um Protein G Yellow Fluorescent Particles (PGFP-0552-5, Spherotech) at 4°C for 1h. SiRPα-Fc-coated beads were subsequently incubated with tumor cells at 4°C for 1h and cell binding was assessed by detecting the fluorescent beads. Binding of T cell-secreted SiRPα-Fc on tumor cell CD47 was assessed by incubation of tumor cells with 24h-collected transduced CD8^+^ or CD4^+^ T cell culture supernatants, followed by staining with labeled goat anti-human IgG Fc (HP6017, Biolegend) at 4°C for 30min. T cell transduction with constitutive SiRPα-Fc, SiRPα monomers, A4-Fc and Fc monomers was detected by eGFP expression. T cell transduction with inducible SiRPα-Fc and SiRPα monomers was detected by eGFP expression, after 48h stimulation of the cells with PMA/Ionomycin (Cell stimulation Cocktail, Thermo Fisher Scientific). Binding of T cell-secreted SiRPα-Fc on T cell CD47 was assessed by staining with labeled goat anti-human IgG Fc (HP6017, Biolegend) at 4°C for 30min. Binding of T cell-secreted A4-Fc on OT-I CD47 was assessed by staining with labeled goat anti-mouse IgG Fc (polyclonal, antibodies-online) at 4°C for 30min. In the absence of a tag, binding of T cell-secreted SiRPα monomer on T cell CD47 was assessed indirectly by staining with anti-human CD47 (CC2C6, Biolegend) at 4°C for 30min, whereas binding of T cell-secreted A4 monomer on OT-I T cell CD47 was assessed indirectly by staining with anti-mouse CD47 (miap301, Biolegend) at 4°C for 30min.

### TCR detection on transduced T cells

Expression of TCRs on the surface of transduced T-cells was detected either by specific fluorescently conjugated multimers from the Tetramer Core Facility at the University of Lausanne, or by staining with anti-human TCR mAbs (Vβ13.1-IMMU 222, Beckman Coulter for A2/NY A97L-TCR). Cell staining was performed at 4°C for 30min.

### Intranuclear Ki-67 detection in T cells

For detection of intranuclear Ki-67, cells were fixed and permeabilized with the FoxP3 transcription factor staining buffer set (Thermo Fisher Scientific) at 4°C for 1h, and subsequently stained with an anti-human Ki-67 (Ki-67, Biolegend) mAb at RT in the dark for 45min. DAPI (Sigma) or Live/Dead fixable Aqua Dead (Thermo Fisher Scientific) cell staining were used to exclude dead cells, according to the manufacturer’s instructions. Apoptotic cells were excluded by staining with Annexin V (BD Biosciences) at 4°C in the dark for 15min. Cell acquisition was performed on a LSRII flow cytometer (BD Biosciences) and data were analyzed using FlowJo (TreeStar).

### Production of retrovirus and lentivirus

To produce lentiviral particles HEK293T cells were co-transfected with 15ug pRRL transfer plasmid, and 7ug pVSV-G and 18ug R874 (containing Rev and Gag/Pol) lentiviral packaging plasmids. For the production of retroviral particles HEK293T cells were co-transfected with 21ug pMSGV transfer plasmid and 18ug pMD22.-Gag/Pol and 7ug pMD RD114 (feline endogenous virus envelope glycoprotein) retroviral packaging, using a mix of Optimem medium (Thermo Fisher Scientific) and Turbofect (Thermo Fisher Scientific). To produce mouse ecotropic retroviral particles, Phoenix-ECO cells were co-transfected with 21ug pMSGV transfer plasmid and 14ug pCL-ECO retroviral packaging plasmid. Culture supernatants were collected 48h or/and 72h post-transfection and concentrated by ultracentrifugation at 24.000×g for 2 hours. Concentrated virus was stored at -80°C until use. Viral titers and MOIs were determined by eGFP reporter gene expression in HEK293T cells.

### Human T-cell isolation, stimulation, viral transduction, and expansion

Healthy donor buffy coat products were purchased from the Transfusion Interrégionale CRS SA (Epalinges, Switzerland) with written consent under a University Institutional Review Board approved protocol. PBMCs were prepared using Lymphoprep (Axis-Shield) density gradient centrifugation and CD8^+^ or CD4^+^ T cells were isolated using CD8 or CD4 magnetic Microbeads (Miltenyi), following the manufacturer’s instructions. Isolated CD8^+^ or CD4^+^ T cells were stimulated with anti-CD3/CD28 beads (Thermo Fisher Scientific) at a 2:1 bead: T cells ratio in the presence of 50IU/ml human IL-2 (GlaxoSmithKline). Lentiviral transduction of activated T cells was performed 24h post-activation by direct addition of the viral particles in the culture medium (MOI 20) and was enhanced by concurrent addition of Lentiboost following the manufacturer’s recommendation (Sirion Biotech). Retroviral transduction of T cells was performed 48h post-activation. Briefly, T cells were transferred to retronectin-coated (Takara) plates previously spinoculated with retroviral particles at 2000×g for 1.5h. T cells were removed from retronectin-coated plates the next day. CD3/CD28 beads were removed when indicated, 3- or 5-days post-activation, and the T cells were maintained thereafter in RPMI 1640-Glutamax (Thermo Fisher Scientific) supplemented with 10% heat-inactivated FCS, 1% Penicillin/Streptomycin, 10ng/ml human IL-7 (Miltenyi) and 10ng/ml IL-15 (Miltenyi) at 0.5-1×10^6^ T cells/ml until downstream use. Live T cells were typically counted manually every 2-3 days, using 0.1% Trypan Blue Stain (Thermo Fisher Scientific) to determine viability, and absolute cell numbers were calculated. Phenotypic analysis of the cells for transduction efficiency was performed after day 7 of culture. For NFAT reactivation, T cells were stimulated for 48h with PMA/Ionomycin (Cell stimulation Cocktail, Thermo Fisher Scientific) or PHA (Sigma).

### Murine OT-I T cell isolation, stimulation, viral transduction, and expansion

OT-I T cells were isolated from the spleens of OT-I CD45.1^+^ mice, using the mouse T cell isolation kit (StemCell Technologies), and following the manufacturer’s instructions. Isolated T cells were stimulated with anti-CD3/CD28 beads (Thermo Fisher Scientific) at a 2:1 bead: T cells ratio in the presence of 50IU/ml human IL-2 (GlaxoSmithKline). Retroviral transduction of T cells was performed 48h post-activation as previously described (23). Briefly, murine T cells were transferred to retronectin-coated (Takara) plates previously spinoculated with retroviral particles at 2000×g for 1.5h. T cells were removed from retronectin-coated plates the next day. CD3/CD28 beads were removed 7-days post-activation and the T cells were maintained thereafter in complete medium supplemented with 10% heat-inactivated FCS, 1% Penicillin/Streptomycin, 10ng/ml human IL-7 (Miltenyi) and 10ng/ml IL-15 (Miltenyi) at 0.5-1×10^6^ T cells/ml until downstream use. Phenotypic analysis of the cells for transduction efficiency was performed after day 7 of culture.

### IFNγ production and T cell cytotoxicity assays by flow cytometry

For the assays, 10^5^ rested NY-ESO-1 TCR^+^ T cells (4:1 CD8^+^:CD4^+^) were co-cultured with 10^5^ NucLight red^+^ tumor cells in complete medium for 24-48h. T cell numbers were normalized based on transduction efficiency and non-transduced T cells were added when needed to achieve similar T cell frequencies among conditions. IFNγ levels in collected cell-free culture supernatants were determined by ELISA (Thermo Fisher Scientific), following the manufacturer’s protocol. T cell cytotoxicity was determined by flow cytometry analysis of the cells and was defined either as the percentage of Annexin V^+^ / DAPI^+^ tumor cells. Results were normalized to the percentage of Annexin V^+^ / DAPI^+^ in cultures of tumor cells alone.

### T cell cytotoxicity assay by IncuCyte live-cell imaging

For the assay, 1.5×10^4^ rested NY-ESO-1 TCR^+^ T cells (4:1 CD8^+^:CD4^+^) were co-cultured with 1.5×10^4^ NucLight red^+^ tumor cells in complete medium for up to 96h. T cell numbers were normalized based on transduction efficiency and non-transduced T cells were added when needed to achieve similar T cell frequencies among conditions. Phase and red fluorescence images were acquired every 2h using IncuCyte^®^ ZOOM (Essen Biosciences). Tumor cell growth was determined by following Total Red Object Area / mm^2^ over time.

### T cell proliferation assay

For the assay, 10^5^ rested NY-ESO-1 TCR^+^ T cells (4:1 CD8^+^:CD4^+^) were labeled with 0.5uM CSFE (Thermo Fisher Scientific) and co-cultured with 10^5^ NucLight red^+^ tumor cells in complete medium. T cell numbers were normalized based on transduction efficiency and non-transduced T cells were added when needed to achieve similar T cell frequencies among conditions. T cell proliferation was assessed 5 days later by flow cytometry and was defined as the percentage of live CFSE^-^ T cells in the culture. T cells activated with PMA-Ionomycin (Thermo Fisher Scientific) were used as positive control.

### Production of human SiRPα-Fc molecules by human T cells

CD8^+^ T cells and CD4^+^ T cells were cultured separately in serum-free medium supplemented with 10ng/ml IL-7/IL-15 at a concentration of 10^6^ SiRPα-transduced T cells/ml for 24h, 48h, 72h and 96h. T cell numbers were normalized based on transduction efficiency and non-transduced T cells were added when needed to achieve similar T cell frequencies among conditions. Human SiRPα-Fc levels in collected cell-free supernatants were determined by an in-house developed ELISA.

Briefly, culture supernatants were incubated with immobilized human CD47 (ACRObiosystems). Bound SiRPα-Fc was detected with a secondary biotinylated rabbit anti-IgG Fc monoclonal antibody (Thermo Fisher Scientific) followed by HRP-conjugated streptavidin (Thermo Fisher Scientific). When indicated, soluble CV1 SiRPα-Fc of known concentration was used as standard.

### Generation of human CD14^+^ monocyte-derived macrophages (MDMs)

PBMCs were prepared from healthy donor buffy coat products using Lymphoprep density gradient centrifugation, as described above. CD14^+^ monocytes were positively isolated using CD14 magnetic Microbeads (Miltenyi), following the manufacturer’s protocol. Macrophages were generated by culturing isolated CD14^+^ monocytes in RPMI supplemented with 10% FCS, 1% Penicillin/Streptomycin and 50ng/ml human M-CSF (ImmunoTools) for 7 days and harvesting the adherent fraction. Culture medium was refreshed at days 3 and 6.

### Generation of mouse bone marrow-derived macrophages (BMDMs)

Whole bone marrow cells were isolated by flushing the femurs and tibiae of NSG mice. Macrophages were generated by incubating whole bone marrow cells in DMEM+Glutamax (Thermo Fisher Scientific) medium supplemented with 10% FCS, 1% Penicillin/Streptomycin, 50uM β-mercaptoethanol and 50 ng/mL mouse M-CSF (ImmunoTools) for 7 days and harvesting the adherent fraction. Culture medium was refreshed at days 3 and 6.

### *In vitro* antibody-dependent cellular phagocytosis (ADCP) assays

For the assays, 5×10^4^ human MDMs or murine (NSG) BMDMs were co-cultured with 5×10^4^ PKH26-labeled (Sigma) tumor cells in serum-free medium in 96-well ultra-low adherent plates (Corning) and incubated for 4-6h at 37°C. To test human T cell-derived SiRPα-Fc and SiRPα monomer molecules, either 5×10^4^ transduced T cells were added to the macrophage/tumor co-culture, or the macrophage/tumor culture was performed with 100μl supernatant of decoy engineered T cells (24h post-transduction). As indicated, recombinant human SiRPα-IgG1 Fc fusion proteins, anti-CSPG4-hIgG1 Fc, Cetuximab (Erbitux, Merck), Trastuzumab (Herceptin, Roche), Avelumab (Bavencio, Merck) or matched isotype were added at 10ug/ml. At the end of the co-culture the cells were washed twice and incubated with human or mouse Fc receptor blocking antibodies (BD Biosciences). Human macrophages were stained with anti-CD11b (M1/70) and anti-CD64 (10.1) mAbs (Biolegend). Murine macrophages were stained with anti-CD11b (M1/70) and anti-F4/80 mAbs (Biolegend). Percentage of phagocytosis was calculated as the percentage of PKH26^+^ cells within human CD11b^+^CD64^+^ and mouse CD11b^+^ F4/80^+^ macrophages, respectively.

### Amnis imaging of tumor cell phagocytosis

For the phagocytosis assay, 5×10^4^ human MDMs or murine (NSG) BMDMs were co-cultured with 5×10^4^ PKH26-labeled (Sigma) tumor cells in the presence of the different recombinant SiRPα decoys at 10ug/ml in serum-free medium in 96-well ultra-low adherent plates (Corning) and incubated for 4-6h at 37°C. Samples were run in the ImageStreamX multispectral imaging flow cytometer (Luminex Corporation) and images of 15,000 events were acquired per sample. Cells were analyzed using a 405 nm laser (15mW) for DAPI excitation, a 488 nm laser (100mW) for PKH26 dye excitation and a 642 nm laser (130 mW) for APC excitation. Brightfield, side scatter and fluorescent cell images were acquired at 40X magnification. Each experimental file contained imagery for 10,000 cells with each cell analyzed for brightfield, side scatter (SSC, Ch06) and three fluorescence channels (DAPI nuclear stain, PKH26 dye and anti-human CD11b-APC). Single color controls were acquired to generate a compensation matrix that was applied to all the experimental files prior to analysis using IDEAS 6.2 software. Only events with APC areas greater than 60 µm² to exclude cell debris and non-saturating pixels were collected as described (85). Images of DAPI^-^ APC^+^ PKH26^+^ double positive events were analyzed downstream with IDEAS 6.2 software. To determine phagocytosis of tumor cells by macrophages, we measured the distance between the center of the PKH26 and APC images of DAPI^-^ APC^+^ PKH26^+^ double positive cells using the Delta Centroid XY (DC) feature. Events in which macrophages have phagocytosed target/tumor cells have lower DC values as compared to aggregate events.

### *In vivo* Winn assay

NSG mice aged 6-12 weeks were subcutaneously inoculated on the flank with 3×10^6^ Saos-2 or Me275 tumor cells previously mixed with 6×10^6^ (or otherwise indicated) A2/NY TCR-T cells coengineered or not to secrete different SiRPα decoys as indicated (4:1 CD8^+^:CD4^+^). Or, mice were similarly subcutaneously inoculated with A2/NY TCR T cells and tumor cells but then administered 1mg Cetuximab and/or 0.2mg Avelumab intraperitoneally twice per week. Start of treatment coincided with T cell injection and continued throughout the duration of the experiment, unless otherwise indicated. Tumor growth was monitored by caliper measurements twice or thrice per week and tumor volume was calculated using the formula V = ½ (L × W^2^) where L is the greatest longitudinal diameter and width is the greatest transverse diameter. Mice were sacrificed when tumors reached 1000mm^3^, lost >20% of original weight or became weak and moribund. Each group consisted of ::5 mice.

### T-cell transfer in xenograft and syngeneic tumor models

6-12 weeks-old female NSG mice were subcutaneously inoculated on the flank with 10^6^ Saos-2 osteosarcoma or A375 melanoma cells. 6-12 weeks-old male NSG mice were subcutaneously inoculated on the flank with 5×10^6^ Me275 cells. Concurrently, human T cells were activated, transduced, and expanded as described above. T cells were adoptively transferred to mice when tumors reached 50-100mm^3^. Mice were treated twice with 10^7^ TCR-expressing SiRPα-Fc- or SiRPα monomer-secreting T cells (4:1 CD8^+^:CD4^+^) or equivalent number of untransduced (NT) or mock-transduced T cells, with the second ACT performed 2-3 days after the first one. Where indicated, 1mg Cetuximab and/or 0.2mg Avelumab mAbs were administered intraperitoneally twice per week. Cetuximab and Avelumab F(ab’)2 fragments were generated using the Pierce F(ab’)2 preparation kit (Thermo Fisher Scientific), following the manufacturer’s instructions, and were administered intraperitoneally twice per week at equimolar amounts as their intact mAb counterparts. Start of mAb and F(ab’)2 treatment coincided with the first T cell transfer and continued throughout the duration of the experiment, unless otherwise indicated.

Female C57BL/6 mice aged 6-12 weeks were subcutaneously inoculated on the flank with 10^5^ B16-OVA melanoma cells. Concurrently, OT-I T cells were activated, transduced, and expanded as described above. T cells were intravenously transferred to mice when tumors reached 50-100mm^3^. Mice were treated twice with 5×10^6^ OT-I A4-Fc- or A4 monomer-secreting T cells, or equivalent number of untransduced (NT) or mock-transduced T cells, with the second ACT performed 2-3 days after the first one.

Tumor growth was monitored by caliper measurements twice or thrice per week and tumor volume was calculated using the formula V = ½ (L × W^2^) where L is the greatest longitudinal diameter and width is the greatest transverse diameter. Mice were sacrificed when tumors reached 1000mm^3^, lost >20% of original weight or became weak and moribund. Each group consisted of ::5 mice.

### Ex vivo studies

To characterize in vivo responses to treatment, mouse tissues were collected at endpoint as indicated. All freshly harvested organs were weighed before dissociation. Tumors and lungs were excised, minced using a scalpel, and enzymatically dissociated in Liberase (Roche) at 37°C for 1h. Single cell suspensions were prepared by mechanical dissociation over a 70μm strainer (Greiner). Harvested spleens were mechanically dissociated over a 40μm strainer (Greiner). Blood samples were collected via cardiac puncture just prior to sacrifice. Single-cell suspensions were depleted of red blood cells (Qiagen) before downstream use.

### Antibodies used for flow cytometric analysis

| RESOURCE | Clone | Source | Identifier | Concentration |
| --- | --- | --- | --- | --- |
| Antibodies for Flow Cytometry |  |  |  | µl per 100µl or working concentration / dilution |
| Human CD4-Phycoerythrin (PE) | SK3 | Biolegend | #344606 | 0.5ul |
| Human CD4-Allophycocyanin (APC) | SK3 | Biolegend | #344614 | 1ul |
| Human CD4-Brilliant Violet 605 (BV605) | OKT4 | Biolegend | #317438 | 0.5ul |
| Human CD4-Spark UV 387 (BUV387) | SK3 | Biolegend | #344686 | 2ul |
| Human CD8-Fluorescein Isothiocyanate (FITC) | SK1 | Biolegend | #344704 | 1ul |
| Human CD8- Allophycocyanin (APC) | SK1 | Biolegend | #344722 | 1ul |
| Human CD8-Brilliant Violet 650 (BV650) | RPA-T8 | Biolegend | #301042 | 0.5ul |
| Human/Mouse CD11b Phycoerythrin-Cy7 (PE-Cy7) | M1/70 | Biolegend | #101215 | 1ul |
| Human CD45-Phycoerythrin-Cy5 (PE-Cy5) | H130 | Biolegend | #304010 | 0.1ul |
| Human CD45-Brilliant Violet 785 (BV785) | HI30 | Biolegend | #304048 | 0.5ul |
| Human CD45RA-ECD | 2H4 | Beckman Coulter | #B49193 | 0.5ul |
| Human CD47-Fluorescein Isothiocyanate (FITC) | CC2C6 | Biolegend | #323106 | 1ul |
| Human CD47- Phycoerythrin-Cy7 (PE-Cy7) | CC2C6 | Biolegend | #323114 | 1ul |
| Human CD64 (Fc receptor I)- Allophycocyanin | 10.1 | Biolegend | #305014 | 1ul |
| Human CD69-Alexa Fluor 700 (AF700) | FN50 | Biolegend | #310922 | 1ul |
| Human CD137 (4-1BB)-Brilliant Violet 711 (BV711) | 4B4-1 | BD Biosciences | #740798 | 1ul |
| Human CD197 (CCR7)- Allophycocyanin-Cy7 (APC-Cy7) | G043H7 | Biolegend | #353212 | 2ul |
| Human CD223 (LAG3)-PerCP-eFluor710 | 3DS223H | Thermo Fisher Scientific | #46-2239-42 | 1ul |
| Human CD274 (PD-L1)- Brilliant Violet 421 (BV421) | 29E.2A3 | Biolegend | #329714 | 1ul |
| Human CD279 (PD-1)- Allophycocyanin-Cy7 (APC-Cy7) | NAT105 | Biolegend | #467415 | 1ul |
| Human CD340 (HER2)-Brilliant Violet 421 (BV421) | 24D2 | Biolegend | #324420 | 2ul |
| Human CD366 (TIM3)-Brilliant Violet 421 (BV421) | F38-2E2 | Biolegend | #345013 | 1ul |
| Human EGFR-Phycoerythrin (PE) | AY13 | Biolegend | #352903 | 2ul in cell pellet |
| Human MCSP (CSPG4)- Fluorescein Isothiocyanate (FITC) | EP-1 | Miltenyi Biotec | #130-098-794 | 1ul |
| Human Ki-67-Brilliant Violet 421 (BV421) | Ki-67 | Biolegend | #350505 | 2ul |
| Human TCR Vbeta13.1- Phycoerythrin (PE) | IMMU 222 | Beckman Coulter | #IM2292 | 7ul in cell pellet |
| Human IgG Fc- Phycoerythrin (PE) | HP6017 | Biolegend | #409304 | 0.5ul |
| Mouse CD11c- Brilliant Violet 605 (BV605) | N418 | Biolegend | #117333 | 1ul |
| Mouse Egr2- Allophycocyanin (APC) | Erongr2 | Invitrogen | #17-6691-82 | 2ul |
| Mouse CD38-Alexa Fluor 488 (AF488) | 90 | Biolegend | #102714 | 2ul |
| Mouse CD45-Phycoerythrin-Cy5 (PE-Cy5) | 30F11 | Thermo Fisher Scientific | #15-0451-82 | 0.2ul |
| Mouse CD45-Phycoerythrin-Cy5.5 (PE-Cy5.5) | 30-F11 | Thermo Fisher Scientific | # 35-0451-82 | 0.1ul |
| Mouse Ter119- Phycoerythrin-Cy7 (PE-Cy7) | Ter119 | Thermo Fisher Scientific | # 25-5921-82 | 0.5ul |
| Mouse F4/80- Allophycocyanin-Cy7 (APC-Cy7) | BM8 | Thermo Fisher Scientific | #47-4801-82 | 1ul |
| Mouse F4/80- Pacific Blue | BM8 | Biolegend | #123124 | 2ul |
| Mouse Ly-6C-PerCP-Cyanine 5.5 (PerCP-Cy5.5) | HK1.4 | Thermo Fisher Scientific | #45-5932-82 | 0.5ul |
| Mouse Ly-6G- Phycoerythrin-Cy7 (PE-Cy7) | 1A8 | Biolegend | #127618 | 1ul |
| Mouse CD172a (SiRPa)- PE-Dazzle | P84 | Biolegend | #144015 | 1ul |
| Annexin V-V500 |  | BD Biosciences | #561501 | 2.5ul |
| Multimer against HLA-A*0201 / NY-ESO-1157-165 SLLMWITQC - Phycoerythrin (PE) |  | TCMetrix / Tetramer Core Facility in University of Lausanne |  | 2ul in cell pellet |
| Multimer against HLA-A*0201 / Melan-A-MART-126-35 EAAGIGILTV - Phycoerythrin (PE) |  | TCMetrix / Tetramer Core Facility in University of Lausanne |  | 1ul in cell pellet |
| Human TruStain FcX™ (Fc Receptor Blocking Solution) |  | Biolegend | #422301 | 10ug/ml |
| Purified Rat Anti-Mouse CD16/CD32 (Mouse Fc Block) | 2.4G2 | BD Pharmingen | #553142 | 10ug/ml |
| Invitrogen™ LIVE/DEAD™ Fixable Aqua Dead Cell Stain Kit, for 405 nm excitation |  | ThermoFisher Scientific | #L34957 | 1/200 |
| DAPI (4',6-Diamidino-2-Phenylindole, Dilactate) |  | Sigma-Aldrich | #D9542 | 100ng/ml |

### Statistics

All statistical analyses were performed on GraphPad Prism 6 software. Statistical tests used for each figure are described in the corresponding figure legend. P values <0.05 were considered statistically significant. Mean ± standard deviation was used to summarize the data, unless noted otherwise. Statistical differences in means of two groups were calculated by two-tailed parametric Student t-tests for unpaired data. Statistical comparisons in means of three groups or more were performed by one-way analysis of variance (ANOVA) or two-way ANOVA with correction for multiple comparisons using Tukey’s test (all groups compared) or Sidak’s test (two select groups compared). The Kaplan-Meier method was used to generate median survival, which was statistically analyzed by log-rank test. No statistical tests were used to predetermine sample sizes.

### Study approval

All in vivo experiments were conducted in accordance with the Service of Consumer and Veterinary Affairs (SCAV) of the Canton of Vaud and Swiss federal law.

### Data availability

Data are available upon request. All supporting data values have been provided as an XLS file to JCI.

## Author contributions

M.I. directed the study. G.C. and R.S. provided advice. E.S. and M.I. conceived and planned the experiments. E.S., A.S., J.P. and B.S. performed the experiments. K.S. provided technical advice. O.M. and V.Z. developed the A2/NY TCR. E.S., M.I., and A.S. analyzed data. E.S., M.I., A.S., and J.P. contributed to the interpretation of the results. E.S. wrote the first draft of the manuscript. M.I. revised the manuscript and together with E.S. finalized it.

## Acknowledgements

We wish to thank all members of the Irving HI-TIDE T-cell engineering group for their technical advice and support over the years, in particular, Greta Giordano-Attianese, Romain Vuillefroy de Silly, Aodrenn Spill, Patrick Reichenbach, Jesus Corria Osorio, Evripidis Lanitis and Catherine Ronet. We wish to thank Dominique Vanhecke for the design and conception of the strategy to modify T cells to express CD47 decoys in the tumor microenvironment, as well as for the initial training and supervision of the research work conducted by E.S. and A.S. We wish to thank Prof. Mikäel Pittet (University of Geneva) for his advice on characterization of the myeloid compartment and careful review of our manuscript. We wish to acknowledge the excellent support of the flow cytometry facility and vivarium at the Epalinges UNIL campus. This work was supported by generous funding from Ludwig Cancer Research, Cancera, the Prostate Cancer Foundation (to G.C.) and the Swiss National Science Foundation (SNF 310030_204326 awarded to M.I., and SNF 31003A_176168 awarded to O.M.).

