## Supplemental figures for "Combination of NY-ESO-1-TCR-T-cells coengineered to secrete SiRPα decoys with anti-tumor antibodies to augment macrophage phagocytosis"

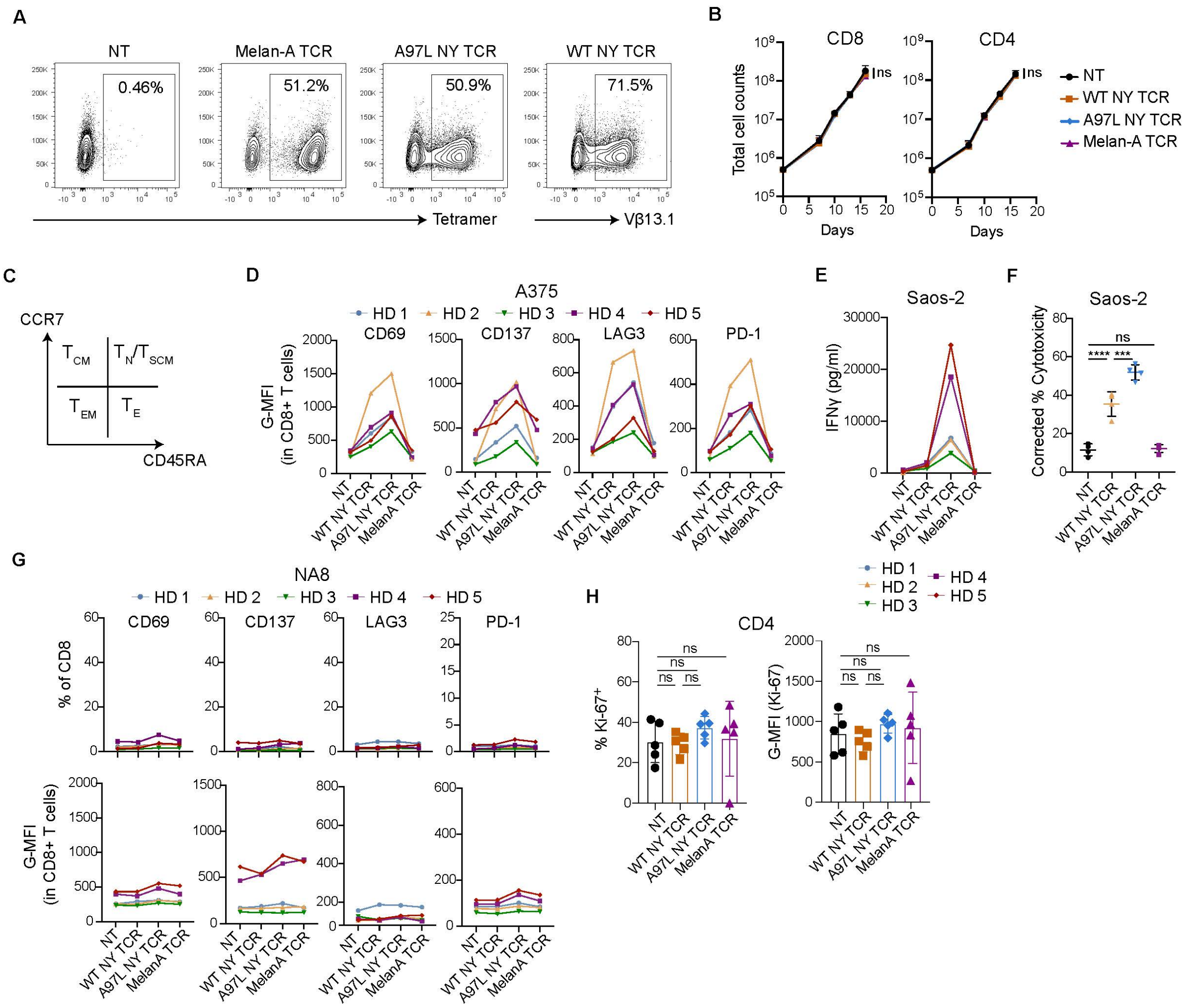


**Supplemental Figure 1. Phenotype and function of TCR-engineered T cells. A)** Transduction efficiency of $CD4^+$ T cells with TCRs evaluated by tetramer or anti-Vβ13.1 mAb staining (data representative of n=5 donors). **B)** Total counts of $CD8^+$ and $CD4^+$ T cells obtained in vitro over 16 days in culture (n=4). **C)** Staining and gating strategy to evaluate effector and memory phenotypes within cultures of rested T cells, transduced or not to express a TCR (Effector, $T_E$; Effector Memory, $T_{EM}$; Central Memory, $T_{CM}$; Naïve/Stem-cell like Memory, $T_{N/SCM}$). **D)** Expression (G-MFI) of activation markers and checkpoint receptors on $CD8^+$ T cells 24h post-stimulation with $A2^+/NY^+$ Saos-2 tumor cells (n=5), HD: Healthy Donor. **E**) IFNγ secretion levels by TCR-modified T cells 24h post-stimulation with Saos-2 tumor cells at E:T = 1:1 (n=5), HD: Healthy Donor. **F)** Frequency of $AnnexinV^+$ $DAPI^+$ cells, corrected to tumor alone, in 24h co-cultures of Saos-2 tumor cells with TCR-modified T cells at E:T=1:1 (n=4). **G**) Expression (frequency and G-MFI) of activation markers and checkpoint receptors on $CD8^+$ T cells 24h post-stimulation with $A2^+/NY^-$ NA8 tumor cells (n=5), HD: Healthy Donor. **H)** Frequency (left) and G-MFI (right) of Ki-67 expression within intratumoral human CD4+ T cells 7 days post-ACT (n=5, data representative of 2 independent experiments). Statistical analysis by two-way analysis of variance

(ANOVA) (B) or one-way ANOVA (F, H) with correction for multiple comparisons by post hoc Tukey's test (B, F and H). ****P< 0.0001; ***P < 0.001; **P < 0.01; *P < 0.05.

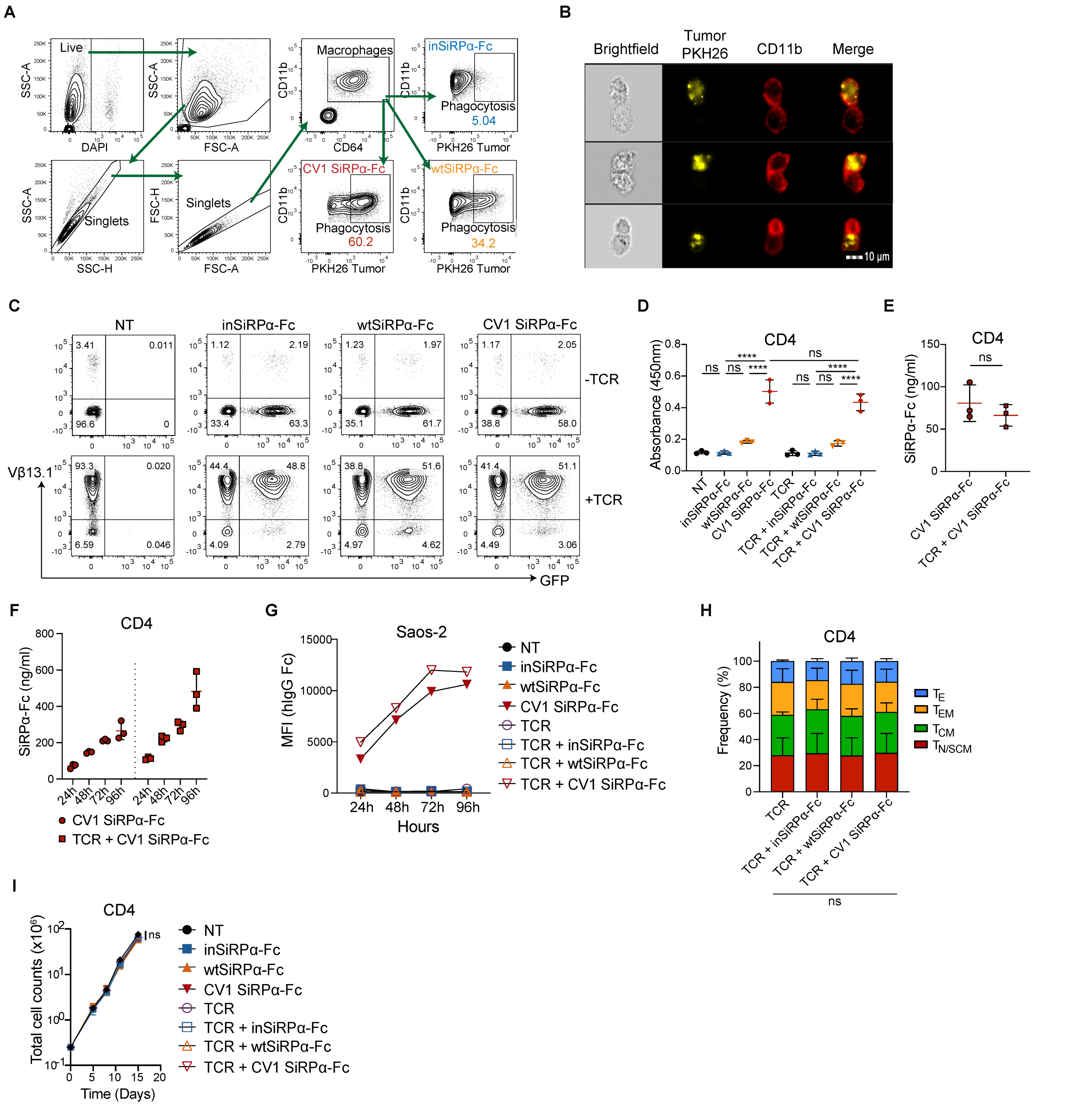


**Supplemental Figure 2. Soluble SiRPα-Fc binds to tumor cell surface CD47 and increases phagocytosis by human macrophages. (A)** Gating strategy for flow cytometric detection of PKH26-labeled tumor cell phagocytosis by human myeloid derived macrophages (MDMs) in vitro. (**B**) Representative Amnis images of human MDMs engulfing PKH26-labeled Saos-2 tumor cells after treatment with 10ug/ml CV1 SiRPα-Fc. **(C)** Expression of SiRPα-Fc and A97L TCR in transduced $CD4^+$ T cells, detected by eGFP and anti-Vβ13.1 mAb staining, respectively (data representative of n=12 donors).

**(D)** CD47-based ELISA detection of SiRPα-Fc secreted by engineered CD4$^+$ T cells (n=3). **(E)** Quantification of CD4$^+$ T cell-secreted CV1 SiRPα-Fc by CD47-based ELISA (n=3). (**F**) Quantification of CV1 SiRPα-Fc accumulated in culture supernatants of engineered CD4$^+$ T cells over time. **(G)** Binding of CD8$^+$ T cell-secreted high-affinity CV1-SiRPα-Fc on Saos-2 using culture supernatants at 24 to 96 hours (data representative of n=2 independent studies). **(H)** Frequency of effector and memory phenotypes of transduced and rested CD4$^+$ T cells (n=3) (Effector, $T_E$; Effector Memory, $T_{EM}$; Central Memory, $T_{CM}$; Naïve/Stem-cell like Memory, $T_{N/SCM}$). **(I)** Expansion of engineered CD4+ T cells (n=3). Statistical analysis by one-way analysis of variance (ANOVA) (D and H), unpaired two-tailed t test (E), or two-way ANOVA (I) with correction for multiple comparisons by post hoc Tukey's test on pooled donors (D and H-I). ****P< 0.0001;***P < 0.001; **P < 0.01; *P < 0.05.

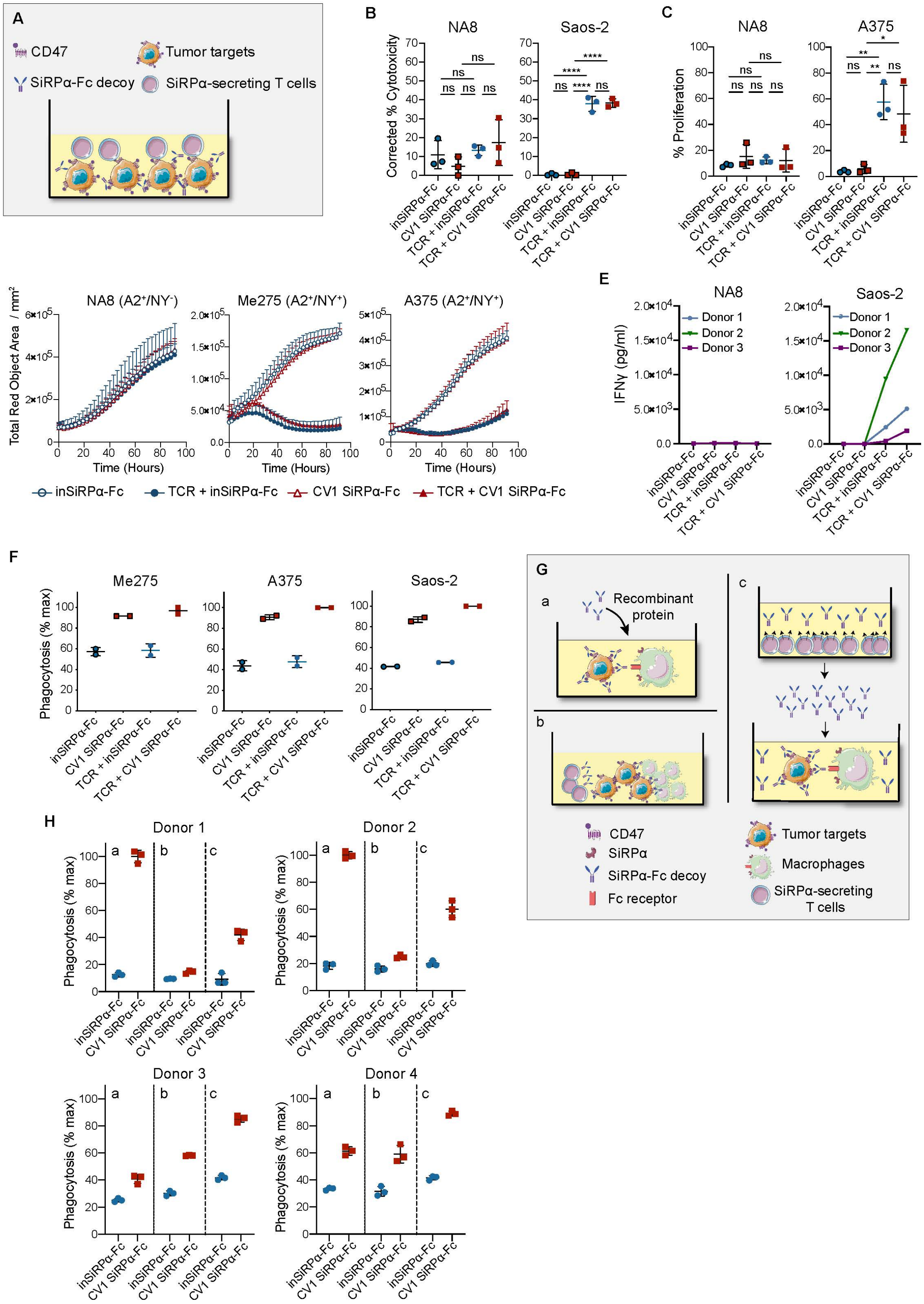
A
CD47
Tumor targets
SiRPα-Fc decoy
SiRPα-secreting T cells
B
NA8
Saos-2
Corrected % Cytotoxicity
ns
****
inSiRPα-Fc
CV1 SiRPα-Fc
TCR + inSiRPα-Fc
TCR + CV1 SiRPα-Fc
C
NA8
A375
% Proliferation
ns
**
*
D
NA8 (A2⁺/NY⁻)
Me275 (A2⁺/NY⁺)
A375 (A2⁺/NY⁺)
Total Red Object Area / mm²
Time (Hours)
inSiRPα-Fc
TCR + inSiRPα-Fc
CV1 SiRPα-Fc
TCR + CV1 SiRPα-Fc
E
NA8
Saos-2
IFNγ (pg/ml)
Donor 1
Donor 2
Donor 3
F
Me275
A375
Saos-2
Phagocytosis (% max)
G
a
Recombinant protein
b
c
CD47
SiRPα
SiRPα-Fc decoy
Fc receptor
Tumor targets
Macrophages
SiRPα-secreting T cells
H
Donor 1
Donor 2
Donor 3
Donor 4
Phagocytosis (% max)
a
b
c
inSiRPα-Fc
CV1 SiRPα-Fc

**Supplemental Figure 3. Evaluation of CV1 SiRPα T cell function as well as macrophage mediated tumor-cell phagocytosis in the presence of CV1 SiRPα decoys in different co-culture conditions. (A)** Schematic of tumor cell and engineered T cell co-culture to evaluate impact of decoys on effector function. **(B)** Frequency of Annexin $V^+$ $DAPI^+$ cells, corrected to tumor alone, 24h post-stimulation with NA8 or Saos-2 tumor cells (n=3). **(C)** Frequency of proliferating engineered $CD8^+$ T cells following stimulation with NA8 or A375 tumor cells (n=3). (**D**) Fluorescent tracking of $mKate2^+$ tumor cells in co-culture with SiRPα decoy coengineered A97L-TCR T cells by live-cell Incucyte imaging (data representative of n=3 donors). (**E**) IFNγ production by engineered T cells 24h post-stimulation with NA8 and Saos-2 tumor cells (n=3 donors). (**F**) Tumor cell phagocytosis by MDMs in the presence of T cell-secreted SiRPα-Fc decoy molecules from single- or TCR dual-transduced T cell cultures (n=2). **(G)** Schematic of different co-culture conditions to evaluate tumor cell phagocytosis by macrophages in the presence of CV1 SiRPα decoy (a = recombinant decoy protein, b = decoy secretion by T cells in a triple co-culture, c = decoy supernatant). **(H)** Side-by-side comparison of tumor cell phagocytosis under conditions shown in Fig. S3G for MDMs (n=4). Statistical analysis by one-way analysis of variance (ANOVA) (B-C) with correction for multiple comparisons by post hoc Tukey's test (B-C). ****$P< 0.0001$; ***$P < 0.001$; **$P < 0.01$; *$P < 0.05$.

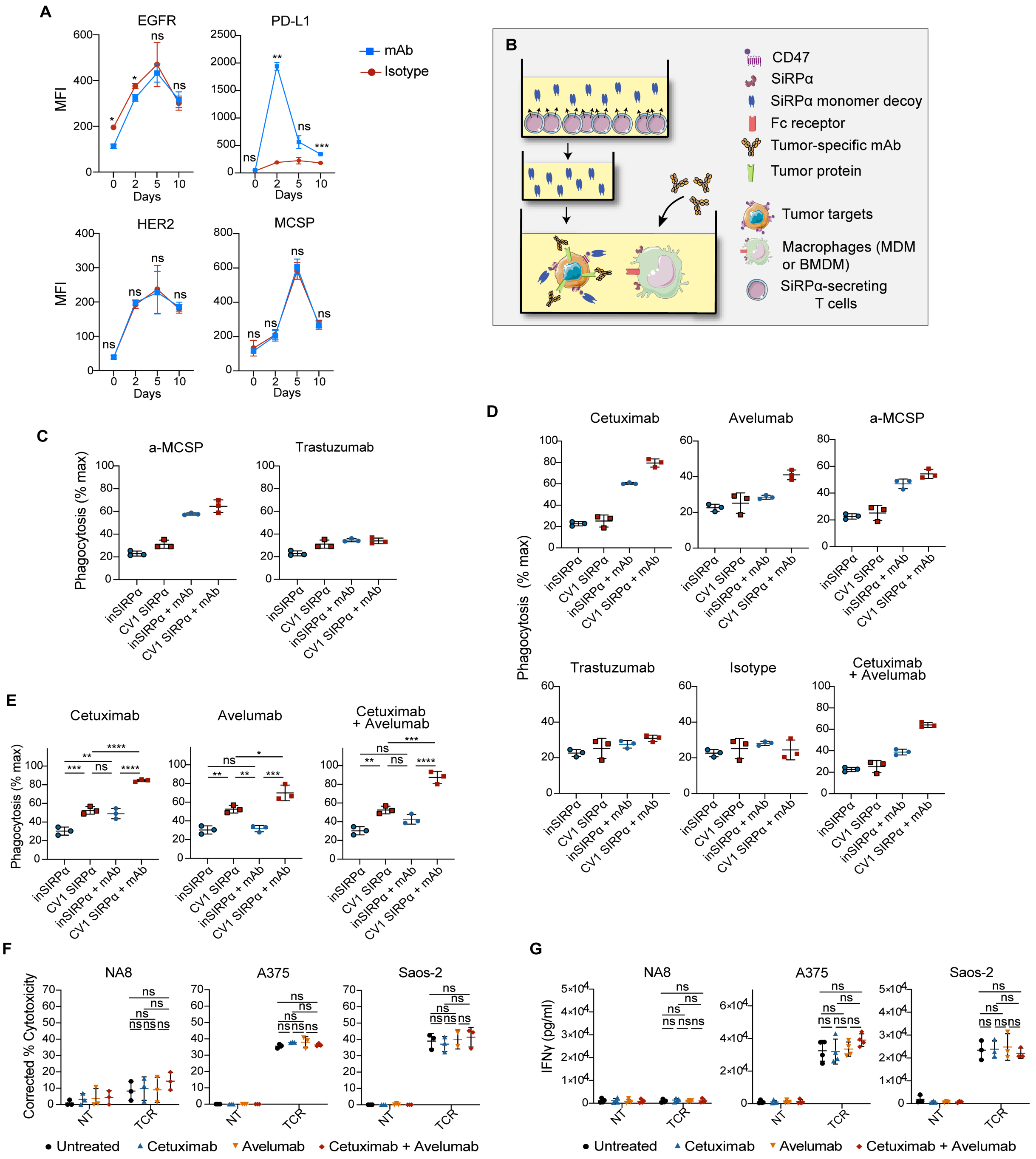

**Supplemental Figure 4. Tumor-targeted monoclonal antibodies synergize with SiRPα monomer secreted by gene-modified T cells to augment macrophage-mediated phagocytosis of tumor cells. A)** Evaluation of EGFR, HER2, MCSP and PD-L1 expression by cultured $CD4^{+}$ T cells at different timepoints (n=3). **(B)** Schematic of macrophage-mediated tumor cell phagocytosis assay in the presence of SiRPα monomer and tumor-targeted mAb. **(C)** Human MDM phagocytosis of A375 tumor cells in the presence of

T cell-secreted SiRPα monomer along with Trastuzumab or anti-MCSP mAbs (representative results for n≥3 donors). **(D)** Human MDM phagocytosis of Saos-2 tumor cells in the presence of T cell-secreted SiRPα monomer along with different tumor-targeted mAbs (representative results for n≥3 donors). **(E)** Murine (NSG) BMDM phagocytosis of Saos-2 tumor cells in the presence of T cell-secreted SiRPα monomer along with Cetuximab or/and Avelumab (n=3). **(F)** Frequency of Annexin $V^+$ $DAPI^+$ cells, corrected to tumor alone, 24h post co-culture with A97L-TCR T cells at E:T = 1:1 in the presence or not of Cetuximab or/and Avelumab (n=3). **(G)** T cell-secreted IFNγ levels 24h post-stimulation with A375 or Saos-2 tumor cells at E:T = 1:1 in the presence or not of Cetuximab or/and Avelumab (n=3). Statistical analysis by two-way analysis of variance (ANOVA) (A) or one-way ANOVA (E-G), with correction for multiple comparisons by post hoc Sidak's test (A) or post hoc Tukey's test (E-G). ****$P< 0.0001$; ***$P < 0.001$; **$P < 0.01$; *$P < 0.05$.

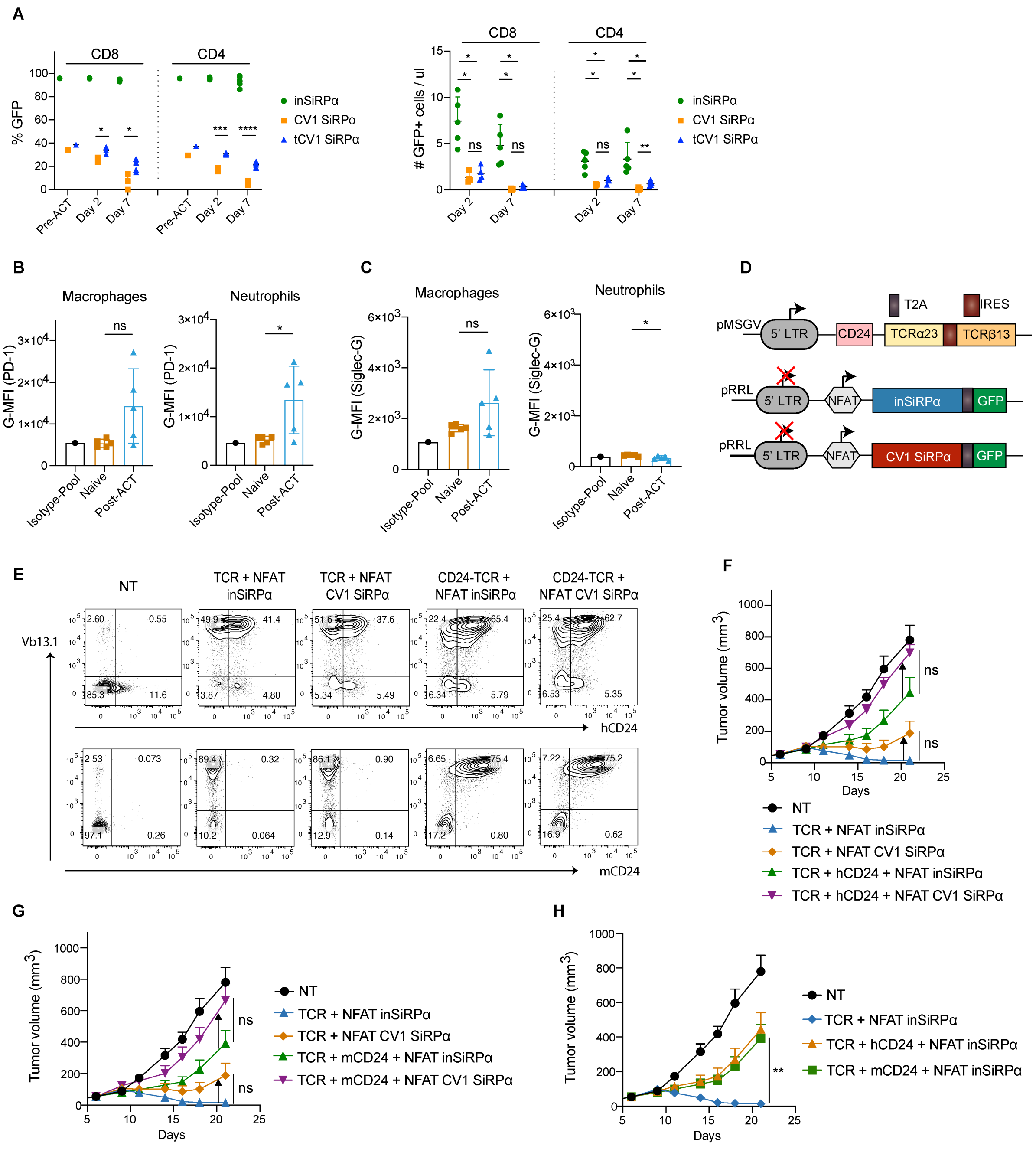


**Supplemental Figure 5. Monomeric SiRPα decoy-engineered human T cells are phagocytosed by NSG-derived macrophages but not by human macrophages. (A)** Frequency (left) and numbers per ul (right) of SiRPα-secreting GFP+ cells in the blood of NSG mice (n≥4, representative results from n=2 independent experiments). **(B)** Expression of mouse PD-1 on the surface of tumor-associated macrophages (left) and neutrophils (left) in established A375 tumors in vivo, pre- and post-ACT (n=5). **(C)** Expression of mouse Siglec-G on the surface of tumor-associated macrophages (left) and neutrophils (left) in established A375 tumors in vivo, pre- and post-ACT (n=5). **(D)** Schematic of lentiviral constructs encoding SiRPα decoys under 6xNFAT and of a retroviral construct encoding mouse or human CD24 together with A97L TCR. **(E)** Expression of human (top panel) or mouse (bottom panel) CD24 and A97L-TCR in transduced $CD8^+$ T cells, detected by anti-CD24 and anti-Vβ13.1 mAb staining, respectively (data representative of n=3 donors). **(F-H)** A375 tumor growth and control curves following ACT with T cells co-engineered with monomeric SiRPα decoys and human CD24 **(F&H)** or mouse **(G&H)** CD24 (n=8, representative results from n=2 independent experiments). Statistical analysis by two-way analysis of variance (ANOVA) (A and F-H) or unpaired, two-tailed t test (B and C) with correction for multiple comparisons by post hoc Tukey's test (A and F-H). ****P< 0.0001; ***P < 0.001; **P < 0.01; *P < 0.05.

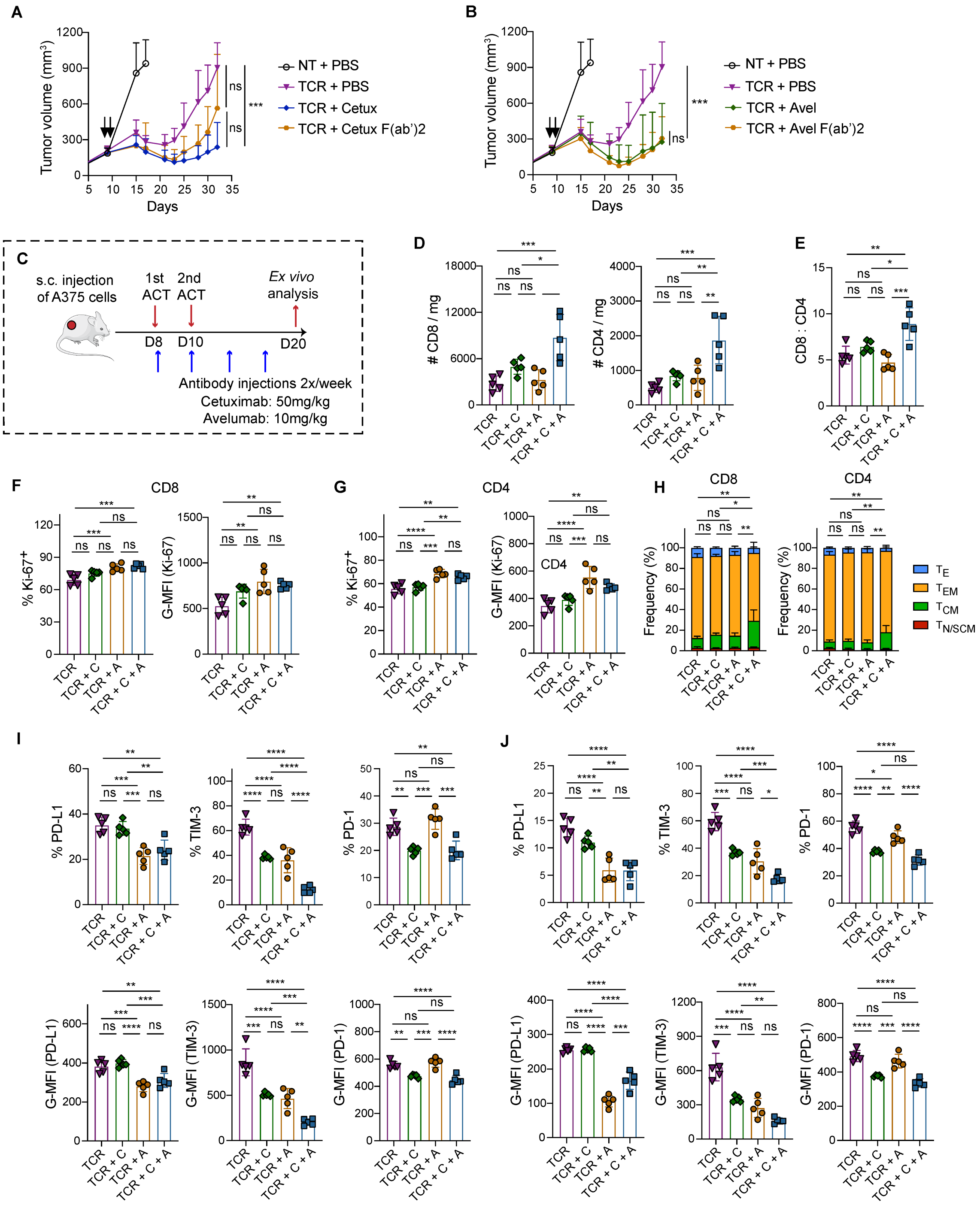

**Supplemental Figure 6. Co-administration of tumor-targeted monoclonal antibodies augments the anti-tumor activity of adoptively transferred A2/NY-TCR T cells. (A)** Control of A375 tumors in NSG mice following adoptive transfer of TCR-modified T cells with co-administration of Cetuximab F(ab')2 fragments or Cetuximab (n=7). **(B)** Control of A375 tumors in NSG mice following adoptive transfer of TCR-modified T cells with co-administration of Avelumab F(ab')2 fragments or Avelumab (n=7). **(C)**

Schematic of ACT study and ex vivo analysis 10 days post-ACT. **(D)** Number of intratumoral human CD8$^{+}$ (left) and CD4$^{+}$ (right) T cells per mg of tumor (n=5). **(E)** Ratio of intratumoral CD8$^{+}$:CD4$^{+}$ human T cell frequency (n=5). **(F)** Frequency (left) and G-MFI (right) of Ki-67 expression within intratumoral human CD8$^{+}$ T cells (n=5). **(G)** Frequency (left) and G-MFI (right) of Ki-67 expression within intratumoral human CD4$^{+}$ T cells (n=5). **(H)** Frequency of effector and memory phenotypes of intratumoral CD8$^{+}$ (left) and CD4$^{+}$ (right) T cells (n=5) (Effector, $T_{E}$; Effector Memory, $T_{EM}$; Central Memory, $T_{CM}$; Naïve/Stem-cell like Memory, $T_{N/SCM}$). **(I)** Frequency (upper panel) and G-MFI (lower panel) of checkpoint receptors in CD8$^{+}$ TCR-engineered T cells (n=5). **(J)** Frequency (upper panel) and G-MFI (lower panel) of checkpoint receptors in CD4$^{+}$ TCR-engineered T cells (n=5). Statistical analysis by two-way analysis of variance (ANOVA) (A-B) or one-way ANOVA (D-J) with correction for multiple comparisons by post hoc Tukey's test (A-B&D-J). ****P< 0.0001; ***P < 0.001; **P < 0.01; *P < 0.05.

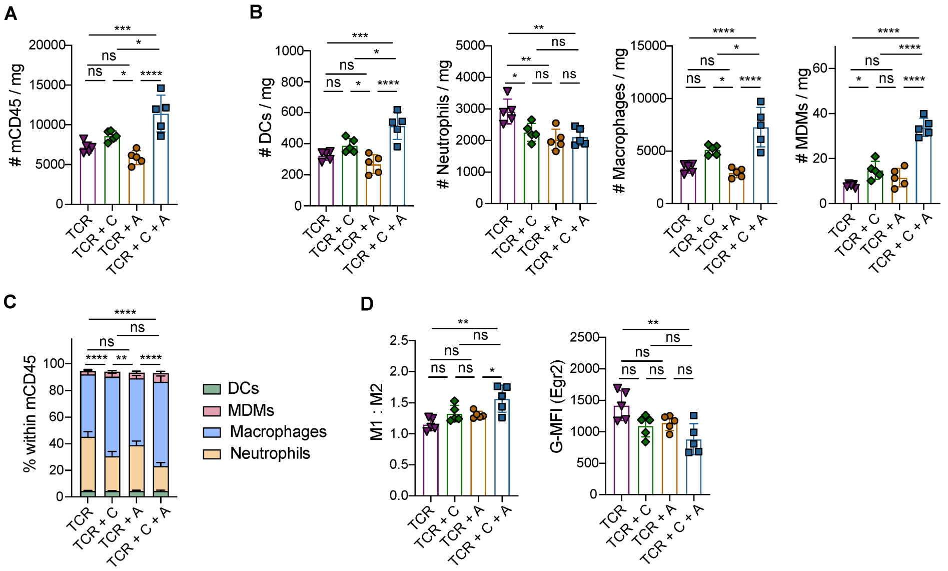


**Supplemental Figure S7. Co-administration of tumor-targeted monoclonal antibodies mobilizes and activates the endogenous innate immune system.** **(A)** Number of intratumoral mouse CD45+ cells per mg of tumor 10 days post-ACT (n=5). **(B)** Number of intratumoral mouse dendritic cells (DCs), neutrophils, macrophages, and monocytic-derived macrophages (from left to right) per mg of tumor (n=5). **(C)** Frequency of mouse myeloid populations within the CD45+ compartment (n=5). (**D)** Ratio of M1:M2 macrophages based on the frequency of CD38+ (M1) and Egr2+ cells (M2) cells, and G-MFI of M2 marker Egr2 (right) (n=5). Statistical analysis by one-way analysis of variance (ANOVA) (A-D) with correction for multiple comparisons by post hoc Tukey's test (A-D). ****P< 0.0001; ***P < 0.001; **P < 0.01; *P < 0.05.
